# Phosphorylation-tuned condensation links HCMV tegument assembly to membrane recruitment

**DOI:** 10.64898/2026.08.05.743017

**Authors:** Yannick Jensen, Laura Cortez Rayas, Boris Bogdanow, Iris Gruska, Barbara Vetter, Enrico Caragliano, Lars Mühlberg, Jens von Einem, Lüder Wiebusch, Jens B. Bosse

**Affiliations:** Hannover Medical School, Institute of Virology, Hannover, Germany; Centre for Structural Systems Biology, Hamburg, Germany; Cluster of Excellence RESIST (EXC 2155), Hannover Medical School, Hannover, Germany; Leibniz Institute of Virology (LIV), Hamburg, Germany; German Center for Infection Research (DZIF), Partner Site Hannover-Braunschweig, Hannover, Germany; Institute of Virology, Ulm University Medical Center, Ulm, Germany; Department of Structural Biology, Leibniz-Forschungsinstitut für Molekulare Pharmakologie (FMP), Berlin, Germany; Institute of Virology, Charité - Universitätsmedizin Berlin, Berlin, Germany; Labor für Pädiatrische Molekularbiologie, Department of Pediatric Oncology and Hematology, Charité - Universitätsmedizin Berlin, Berlin, Germany

## Abstract

Herpesviruses build complex infectious particles around a protein layer, the tegument, that lacks an ordered architecture. How this apparently amorphous material selectively assembles on the capsid and engages enveloping membranes remains unclear. Here we show that the capsid-anchored human cytomegalovirus protein pp150 forms liquid-like condensates when locally concentrated. Its disordered region recruits soluble tegument proteins and membrane-associated partners, providing a mechanism to couple assembly of the tegument layer to recruitment of the enclosing membrane. Phosphorylation tunes this condensation: phosphomimetic mutations prevent recovery of infectious virus, whereas loss of phosphorylation sites causes aberrant capsid–tegument assemblies and impairs viral replication. Together, these findings identify regulated condensation as a mechanism that couples tegument assembly to membrane recruitment and supports the production of infectious particles, highlighting condensate regulation as a potentially novel point of antiviral intervention.

## Introduction

Herpesvirus assembly requires the coordinated construction of a multilayered virion, in which an icosahedral capsid, a surrounding protein layer called the tegument, and a host-derived lipid envelope are brought together with high spatial and temporal precision. The tegument is the dense protein layer that connects the capsid and the outer envelope^1^. After genome packaging in the nucleus, capsids reach the cytoplasm, where they acquire their full tegument and undergo final envelopment by wrapping at virus-induced intracellular membranes^2^. This cytoplasmic phase recruits viral and cellular factors at defined stoichiometries into virions and must interface with the membranes that drive wrapping and release. How herpesviruses organize this capsid-tegument-membrane axis remains poorly understood.

At the center of this problem lies an apparent paradox. In human cytomegalovirus (HCMV), the tegument contains more than 30 viral proteins together with host factors and carries out essential functions during both viral entry and assembly^3–6^, yet it has no obvious ordered architecture^2^. Electron microscopy has long described it as amorphous^7^, and many individual tegument proteins appear partially redundant in cultured cells^8,9^. This combination of compositional complexity, structural plasticity, and functional specificity has made the organizing principles of tegument assembly difficult to define. The central unresolved question is how tegument proteins are selectively concentrated and coupled to the capsids and membranes used for envelopment, whether through stable one-to-one interactions, through recruitment to membranes, or through collective mechanisms that enable many proteins to be concentrated simultaneously.

Liquid-liquid phase separation (LLPS) is one such collective mechanism. Biomolecular condensates form through multivalent, individually weak interactions often mediated by intrinsically disordered regions (IDRs), and they concentrate selected client proteins without a fixed stoichiometry or a rigid scaffold^10–12^. The clients that partition into a condensate are often determined by the amino acid composition and charge patterning of the interacting IDRs rather than by discrete linear motifs^10–12^. Because condensation depends on the balance of these weak interactions, post-translational modification readily tunes it, and phosphorylation is the best-characterized input. Phosphorylation of serine and threonine residues alters the net charge, charge patterning, and interaction valency of an IDR and thereby shifts the concentration at which the protein demixes, so that opposing kinase and phosphatase activities can determine when and where a condensate assembles and dissolves, including in a cell cycle-dependent manner^1^^3,14^. Condensates also interact with lipid bilayers. They wet membrane surfaces and form composite membrane-associated assemblies^15,16^, and in reconstituted and cellular systems, condensate-membrane contacts bend membranes, drive engulfment, and promote scission^17,18^.

Some herpesvirus-induced phase-separated compartments have been described, such as HCMV nuclear replication compartments at viral genomes^19^ or the KSHV ORF52 protein that drives formation of the cytoplasmic assembly compartment through LLPS^20^. Many tegument proteins contain large disordered regions, and for a few, such as UL11 of herpes simplex virus 1 (HSV-1)^21^, they also form condensates in vitro. This suggests that the virion tegument surrounding the capsid may form by condensation. However, it is unclear how and by which viral protein such condensation would be coordinated around the capsid, how interactions between the many tegument proteins and the enveloping membrane is organized and how such complex condensates would be regulated^2^.

The HCMV tegument protein pp150, encoded by UL32, is a strong candidate for this role, as it combines the architecture and the regulatory inputs that such a function requires. Its structured N-terminal region anchors on the capsid surface, where approximately 955 copies form a net-like layer, whereas its disordered C-terminal two-thirds could coordinate tegument condensation^22^. Consistent with this, our previous spatially resolved crosslinking of intact virions identified this IDR as a dominant tegument interaction hub, with contacts spanning capsid-proximal proteins and factors near the envelope^23^. The disordered part of pp150 is also a substrate for both kinases and phosphatases, which might regulate condensation^23,24^. Consistent with these proposed roles, pp150 is essential for viral replication and is one of the most abundant virion components^9,25^.

Here we show that the pp150 IDR indeed forms condensates. Using light-inducible corelet seeding in living cells, we demonstrate that the pp150 IDR undergoes LLPS when clustered at a density similar to what is found on the viral capsid. These condensates dynamically recruit other tegument proteins such as the soluble UL25 and vesicles positive for the membrane-associated tegument protein UL71, illustrating that pp150 condensates can recruit other tegument proteins and engage membranes needed for virion envelopment. Importantly, we show that phosphorylation regulates pp150 condensation. Hyperphosphorylation suppresses condensation, while hypophosphorylation increases it. Recombinant viruses carrying corresponding mutations either yield no infectious progeny or replicate poorly and accumulate aberrant cytoplasmic tegument–capsid assemblies outside the assembly compartment, indicating that phosphorylation restricts pp150 condensation to the correct time and place during infection. Together, these findings identify pp150 as a regulated assembly hub that connects capsid-associated tegument organization with the membrane-proximal machinery of final envelopment. In a related study, Mühlberg et al. identify the HSV-1 tegument protein VP22 as a non-homologous LLPS-competent scaffold that organizes a distributed network spanning capsid-proximal and envelope-associated tegument proteins through an analogous mechanism^26^. Because pp150 (UL32) and VP22 (UL49) are subfamily-restricted proteins without detectable sequence relationship, their convergence on the same organizing principle indicates that condensate-based tegument organization is a general feature of herpesvirus morphogenesis.

## Results

### Clustering the pp150 IDR at capsid-like density drives liquid–liquid phase separation in living cells

Because pp150 is anchored to the capsid through its structured N-terminal domain, its C-terminal IDR is displayed on the capsid surface at a local density far above that of a diffuse cytoplasmic pool, the regime in which weak multivalent interactions can drive phase separation^10^. We therefore asked whether the pp150 IDR forms a liquid condensate when clustered at comparable density in living cells. To test this possibility, we used the corelet system that enables light-induced protein concentration ^27^. In this system, iLID-eGFP-FTH1 forms multivalent 24-mer nanoparticles that act as light-responsive seeds, whereas an SspB-tagged protein of interest is recruited to these particles upon blue-light activation of the iLID-SspB interaction (Fig. 1a, b). If the recruited protein has phase-separation capacity, local concentration can trigger the formation of condensates detectable by fluorescence microscopy.

**Fig. 1:**
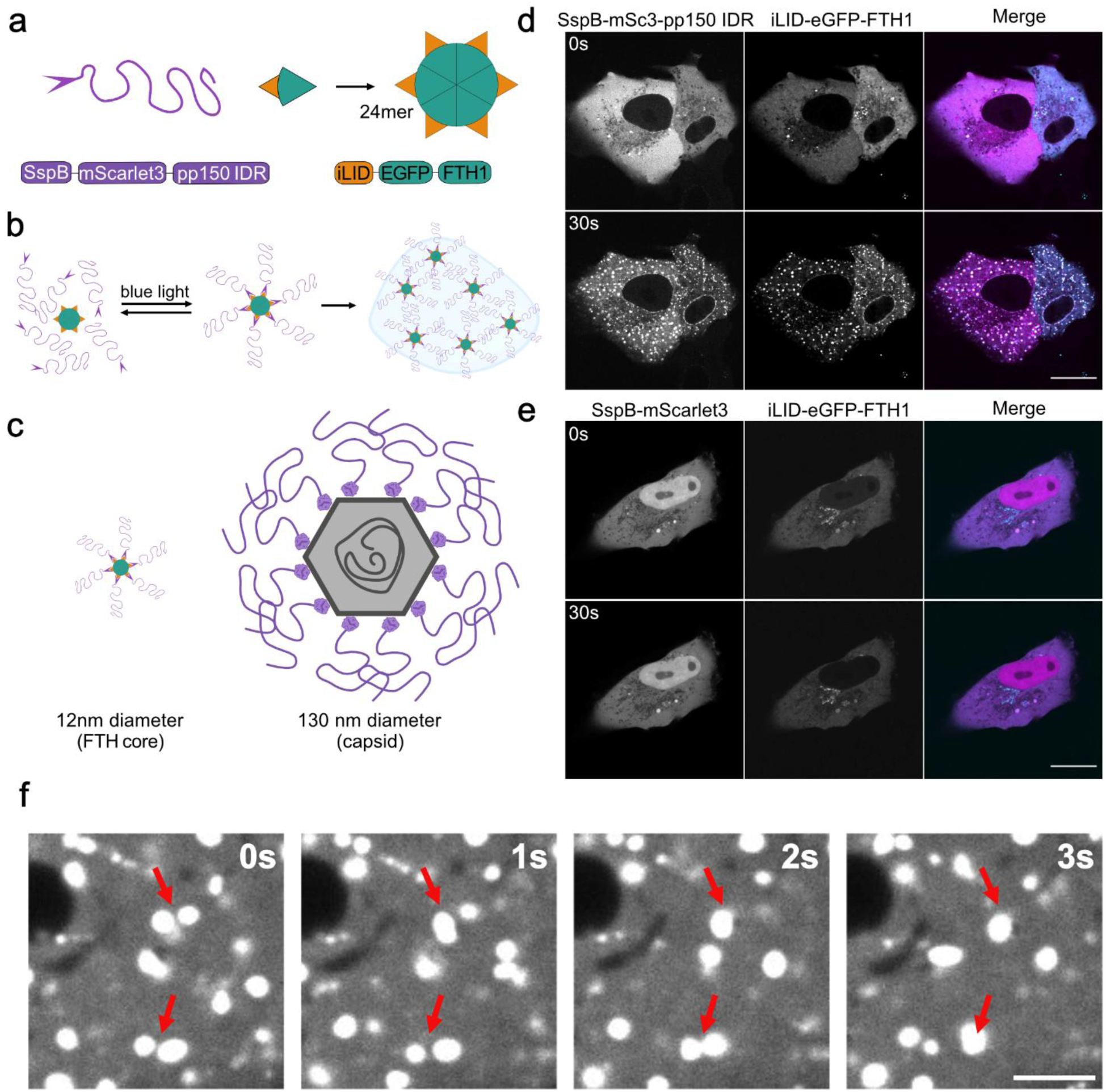
The pp150 IDR forms liquid-like condensates upon local clustering. **a**, Schematic of the SspB-mScarlet3-pp150 IDR construct. **b**, Schematic of the corelet system: iLID-eGFP-FTH1 assembles into 24-mer nanoparticle seeds of approximately 12 nm diameter that recruit SspB-tagged proteins upon blue-light activation, which can trigger condensate formation. **c**, Schematic illustrating that activated pp150 corelets approximate the macromolecular organization of pp150 on the viral capsid. **d**, Live-cell images of VeroB4 cells expressing iLID-eGFP-FTH1 (cyan) and SspB–mScarlet3–pp150 IDR (magenta) before and after blue-light activation. Activation triggers rapid formation of small, round condensates that subsequently fuse, indicating liquid-like behaviour. Scale bar, 20 μm. **e**, As in **d**, for SspB–mScarlet3 lacking the pp150 IDR, which remains diffuse under identical activation conditions. Scale bar, 20 μm. **f**, Magnified time series of fusion events between pp150 IDR condensates. Scale bar, 3 μm. Images in **d**–**f** are representative of three independent transfections performed on separate days (n = 3). For each replicate at least 10 cells were imaged.

To adapt this system to the native organization of pp150 on capsids, we generated an SspB-mScarlet3-pp150^28^ IDR construct, preserving the N-to-C-terminal radial orientation of pp150. Upon corelet activation, the multivalent recruitment of the pp150 IDR to corelet seeds approximates the high local density of pp150 on viral capsids (Fig. 1c). We first probed the intrinsic condensation behavior of the pp150 IDR independently of other viral factors and performed experiments in transfected VeroB4 cells. Before light activation, both SspB-mScarlet3-pp150 IDR and the SspB-mScarlet3 negative control showed a largely diffuse cytoplasmic distribution. Upon blue-light activation, SspB-mScarlet3-pp150 IDR rapidly formed numerous spherical cytoplasmic condensates, whereas SspB-mScarlet3 alone remained diffuse (Fig. 1d, e). The pp150 IDR condensates frequently underwent fusion events and relaxed into round structures, consistent with liquid-like behavior (Fig. 1f).

Together, these data show that the pp150 IDR, when clustered at a density approximating its display on the capsid, is sufficient to undergo inducible LLPS in living cells.

### pp150 condensates recruit other tegument proteins and associate with membranes

Having established that the pp150 IDR can undergo phase separation, we next searched for proteins likely to be recruited into pp150 condensates. We reasoned that if pp150 organizes the tegument through condensate-like interactions, its contacts with partner proteins should be enriched within disordered regions, and that partners contacted through their most disordered segments would be the best candidates for IDR-dependent recruitment. This provides an unbiased criterion for nominating clients, rather than selecting partners by abundance or by prior expectation. Therefore, we reanalyzed our previously published HCMV virions crosslinking dataset^23^. We focused on the lysine residues on partner proteins that crosslink to pp150, and asked whether these contact sites reside in more disordered local environments than expected. Across the testable tegument interactors, lysines crosslinking to pp150 localized to significantly more disordered local regions (predicted with IUPred3^29^) than accessibility-matched background lysines on the same protein (median Δ disorder = +0.065; mean Δ = +0.151; Wilcoxon signed-rank P = 0.0067; 12 of 15 proteins positive, Fig. 2a). For several partners, the pp150-contacting region was not only relatively but absolutely disordered (mean IUPred3 score > 0.5), including UL71, UL97, and UL25 (Fig. 2a). This disorder preference was specific to the tegument. Capsid and capsid-associated proteins, which are also crosslinked to pp150, did not show enrichment of disorder at their contact sites (median Δ = −0.055; Wilcoxon versus zero P = 0.63). Host proteins crosslinking to pp150 likewise lacked enrichment of disorder (median Δ = −0.025; P = 0.21, Supplementary Fig. 1). This analysis indicates that pp150 uses two distinct interaction modes within the same virion, a structured one toward the capsid and a disorder-based one toward the tegument, which is the architecture a capsid-anchored condensate scaffold would require. Based on this analysis, we selected the soluble tegument protein UL25 and the membrane-associated tegument protein UL71 for experimental validation, because both showed high contact-site disorder.

**Fig. 2:**
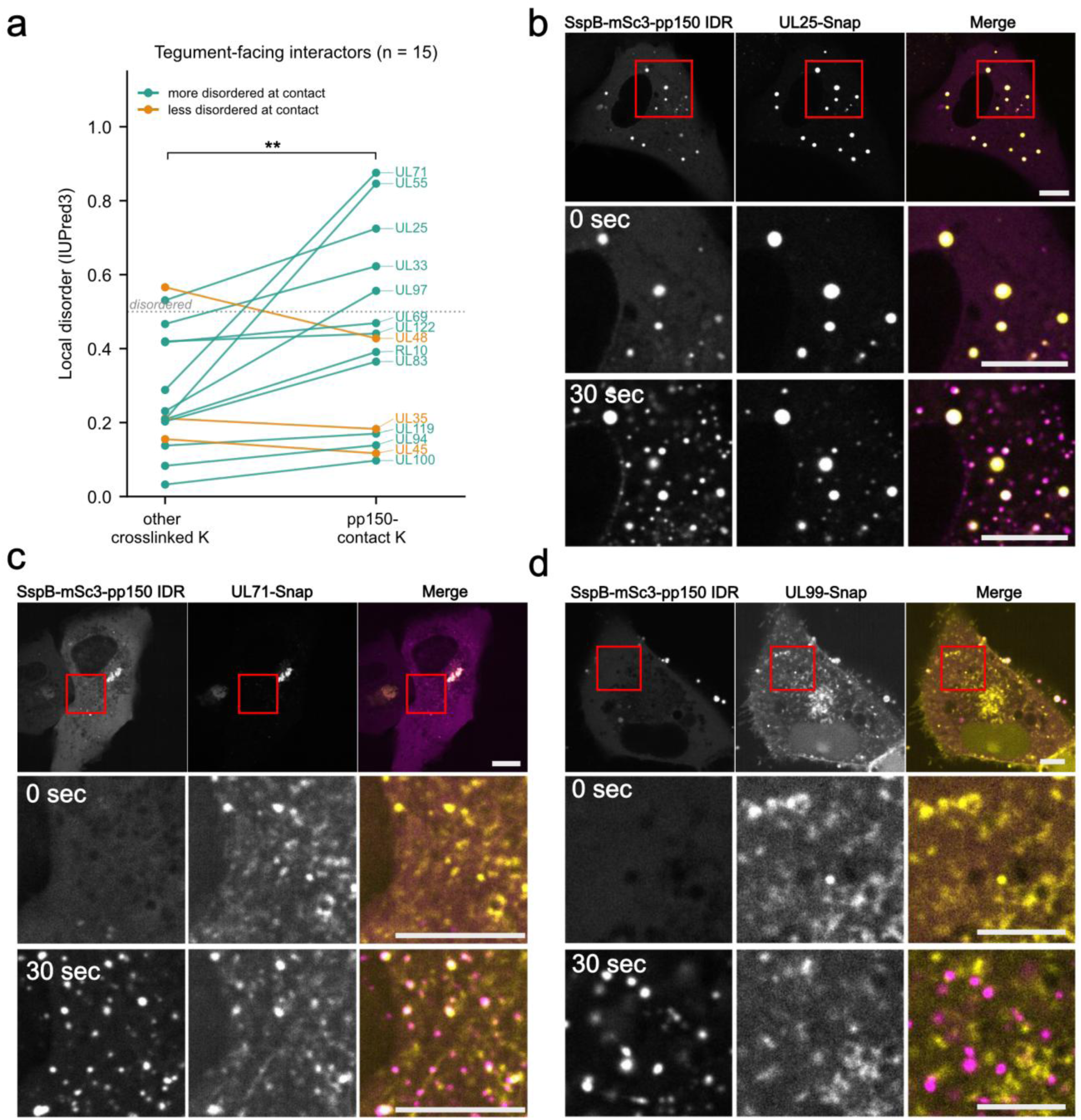
pp150 IDR condensates selectively recruit UL71 and UL25 but not UL99. **a**, Local disorder at pp150 contact sites in tegument-facing virion proteins. For each protein, the mean IUPred3 disorder (long mode) of 15-residue windows (±7 residues, truncated at sequence termini) centred on lysines crosslinked to pp150 was compared with that of windows centred on the remaining crosslinked lysines of the same protein. Windows were averaged within each protein before testing. The experimental unit is the protein, and contact and background values are paired within protein (n = 15 proteins). Mean Δdisorder = +0.151 (s.d. 0.219; 95% CI 0.030 to 0.272), median 0.065; two-sided Wilcoxon signed-rank test, *W* = 14, *P* = 0.0067. The capsid and host contrast groups are shown in Supplementary Fig. 1. **b**, Live-cell images of VeroB4 cells co-expressing the pp150 IDR corelet constructs and UL25–SNAP after blue-light activation. SspB–mScarlet3–pp150 IDR, magenta; UL25–SNAP, yellow. The iLID-eGFP-FTH1 channel is omitted for clarity. Scale bars, 10 µm. **c**, As in b, with UL71–SNAP (yellow). Insets are displayed with a linearly expanded display range (brightness and contrast only) applied uniformly to both channels, because the single bright perinuclear structure in the overview would otherwise saturate the display and render smaller vesicular structures invisible; intensities are therefore not comparable between overview and inset. Scale bars, 10 µm. **d**, As in b, with UL99–SNAP (yellow). UL99 is not recruited into pp150 IDR condensates. Scale bars, 5 µm. Images in b–d are representative of three independent transfections performed on different days (n = 3).

Next, we validated the IDR-dependent recruitment of both proteins to pp150 condensates by co-transfecting SNAP-tagged variants of UL71 or UL25 with the SspB-mScarlet3-pp150 IDR corelet system and imaging condensation behaviour after photoactivation.

Co-expression of UL25 relocalized SspB-mScarlet3-pp150 IDR into discrete UL25-positive structures even before corelet activation and more condensates positive for both proteins appeared upon photoactivation of pp150 condensation, indicating that pp150 condensates can recruit another tegument protein. (Fig. 2b).

Co-expression of UL71-SNAP with the pp150 IDR corelet system relocalized SspB-mScarlet3-pp150 IDR from a diffuse distribution onto small, highly mobile UL71-positive puncta upon corelet activation (Fig. 2c). These puncta were positive for the endocytic tracer FM4-64 FX, showing that UL71 is associated with endosomal vesicles (Supplementary Fig. 2). This suggests that UL71-positive vesicles serve as platforms for pp150 condensation and provides a direct physical link between a pp150-rich tegument phase and intracellular membranes used for envelopment in infection.

We also tested UL99, a membrane-associated tegument protein with extensive disorder. UL99-SNAP was not recruited to light-induced pp150 condensates (Fig. 2d), showing that not all IDR-containing tegument proteins are enriched in these condensates.

### Tegument protein recruitment into condensates depends on IDR amino acid composition rather than sequence order

We next asked whether tegument partners could promote pp150 IDR condensation without optogenetic clustering. Co-expression with either UL71-SNAP or UL25-SNAP induced assemblies containing both the partner and mScarlet3-pp150 IDR, without corelet seeds or blue-light activation. Neither partner recruited mScarlet3 alone, and SNAP alone did not reorganize the pp150 IDR (Fig. 3a, b).

**Fig. 3:**
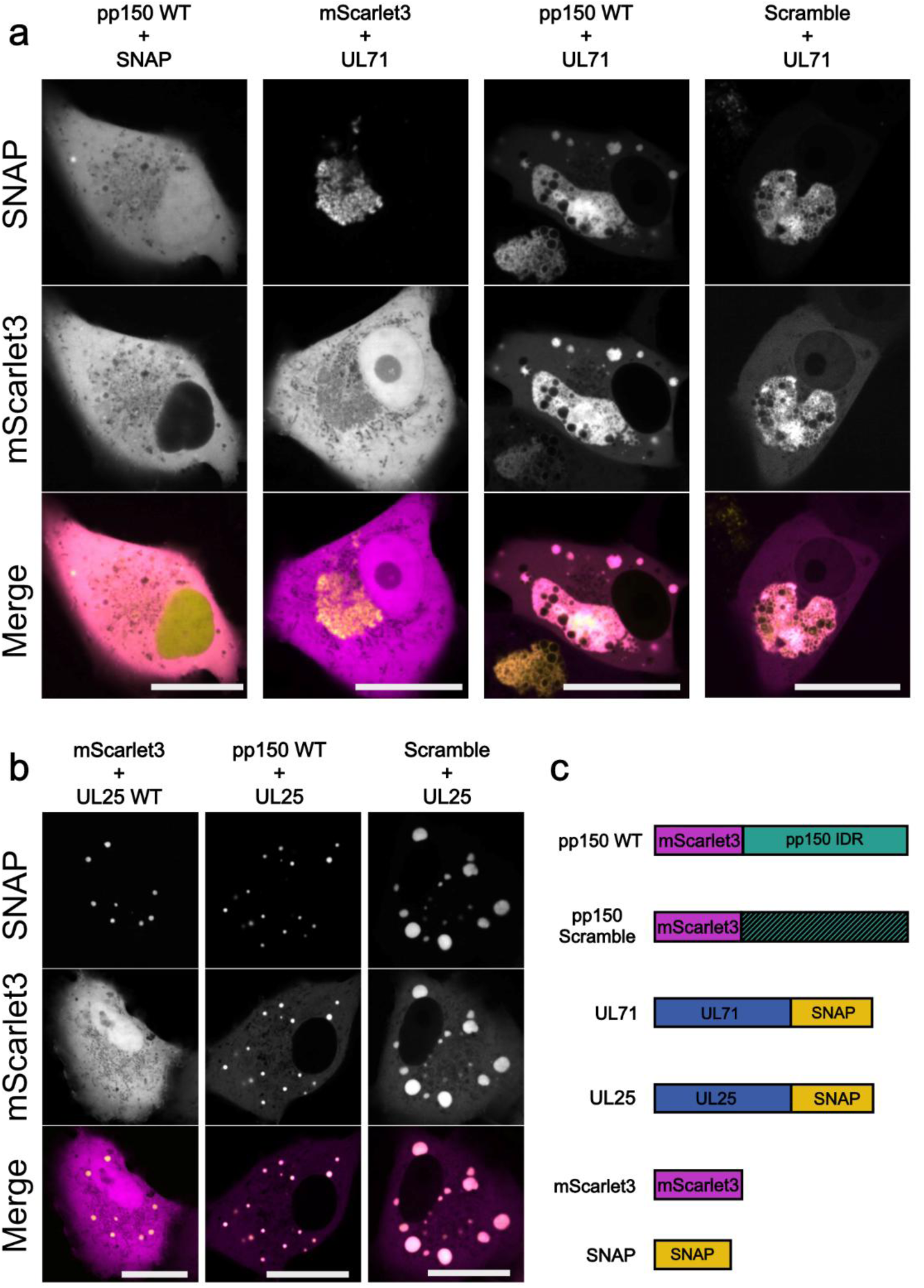
UL71 and UL25 co-condense with the pp150 IDR independently of IDR sequence order. All experiments in this figure were performed without the corelet seed and without blue-light activation. **a**, Live-cell images of VeroB4 cells co-expressing the indicated mScarlet3-pp150 IDR construct (magenta) and UL71-SNAP (yellow). Co-expression with UL71 relocalized mScarlet3-pp150 IDR from a diffuse cytoplasmic distribution into distinct condensate-like accumulations, and scrambling the pp150 IDR sequence did not abolish this phenotype. Controls comprised SNAP alone co-expressed with mScarlet3-pp150 IDR and mScarlet3 alone co-expressed with UL71-SNAP, included to test for tag-driven interactions; neither produced detectable co-condensation or relocalization. Scale bars, 20 µm. **b**, As in a, with UL25-SNAP (yellow). Co-expression with UL25 relocalized mScarlet3-pp150 IDR into distinct UL25-positive condensate-like structures, and scrambling the IDR did not abolish the phenotype. mScarlet3 alone co-expressed with UL25-SNAP produced no detectable co-condensation or relocalization. Scale bars, 20 µm. **c**, Schematic of the pp150, UL71 and UL25 constructs used. Images in a,b are representative of three independent transfections performed on different days (n = 3, at least 10 cells were imaged for each replicate).

These assemblies fused, returned to rounded shapes and showed rapid fluorescence recovery after photobleaching (FRAP). Mobile fractions were 81.7% with UL25 and 80.6% with UL71, compared with 29.6% for the aggregate-forming control MCMV M45-mCherry (P = 0.0023 and P = 0.0011, respectively)^30^. These findings support the formation of dynamic condensates rather than static aggregates (Supplementary Fig. 3).

To test whether co-condensation requires the native pp150 IDR sequence, we scrambled its sequence while preserving its amino acid composition. The scrambled IDR still co-condensed with both UL71-SNAP and UL25-SNAP (Fig. 3a, b). Thus, the native sequence order is dispensable for co-condensation in this assay, supporting a role for amino acid composition in recruitment. Together, these results show that tegument partners can promote pp150 IDR condensation without externally imposed clustering, with membrane-associated UL71 providing a link to membranes.

### Phosphorylation tunes pp150 IDR condensation

If pp150 forms a liquid condensate on the capsid that recruits further tegument partners, it must be regulated to prevent indiscriminate clumping of tegumented capsids. An attractive candidate mechanism is phosphorylation, a well-established regulator of biomolecular condensates^13^. The pp150 IDR is phosphorylated by cyclin-dependent kinases (CDKs)^24^, while protein phosphatase 1 (PP1) is recruited into virions by pp150 and antagonizes tegument protein phosphorylation^23^.

We therefore generated three pp150 IDR corelet variants to test how phosphorylation-related changes affect condensation (Fig. 4a). First, we substituted 32 predicted minimal CDK consensus sites with glutamate (S/T-P→E-P) to mimic constitutive negative charge (hyper-P-mut). Second, we replaced the same residues with alanine (S/T-P→A-P) to generate a phospho-deficient mutant (hypo-P). Third, we included a PP1-binding-deficient mutant (PP1-mut), in which the SILK and RVxF motifs required for PP1 binding were disrupted^23^. Because PP1 binding promotes pp150 dephosphorylation, PP1-mut is expected to increase pp150 phosphorylation and therefore serves as an independent phosphorylation-enhanced condition. Critically, PP1-mut alters only the SILK and RVxF motifs, a small number of residues, and leaves the composition of the IDR otherwise intact, so it separates the consequences of phosphorylation state from those of the extensive sequence changes carried by the hyper-P and hypo-P variants.

**Fig. 4:**
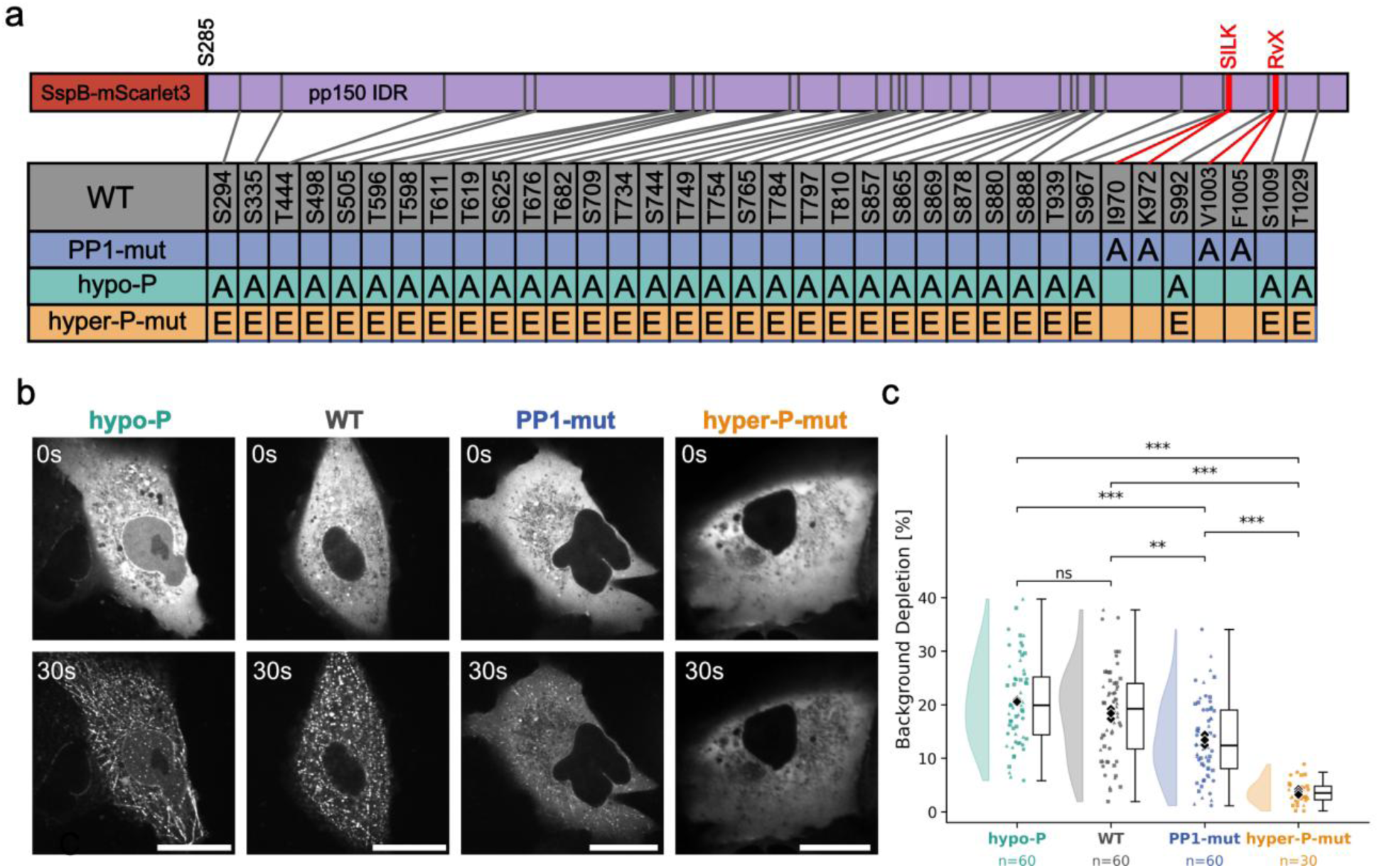
Phosphorylation of the pp150 IDR tunes condensate formation. **a**, Schematic of the pp150 intrinsically disordered region (IDR) showing the 32 proline-directed serine/threonine–proline (S/T-P) sites and the three phospho-variant corelet constructs: substitution of all 32 S/T-P sites with glutamate (hyper-P-mut), a PP1-binding-deficient variant (PP1-mut) and substitution of the 32 cyclin-dependent kinase consensus sites with alanine (hypo-P). **b**, Live-cell images of VeroB4 cells expressing the pp150 phospho-variants in the corelet system before and after activation. hypo-P formed condensates comparable to wild type, PP1-mut formed smaller and less frequent condensates, and hyper-P-mut formed none detectable. Scale bars, 20 µm. Images are representative of three independent experiments. **c**, Condensation efficiency per cell, the percentage reduction in cytoplasmic mScarlet3 fluorescence between the pre-and post-activation frame. Colored squares, triangles and circles, single-cell measurements from biological replicates 1, 2 and 3, respectively; black diamonds, replicate means. Box center line, median; box bounds, 25th and 75th percentiles; whiskers, most extreme data point within 1.5× the interquartile range of the box bounds. *n* = 60 cells (20 per replicate) for wild type, hypo-P and PP1-mut and *n* = 30 cells (10 per replicate) for hyper-P-mut, from three independent transfections. Because hyper-P-mut formed no visible condensates, evenly fluorescent cytoplasmic regions were measured, so its values report the assay noise floor. Comparisons used a linear mixed-effects model with construct as a fixed effect and biological replicate as a random intercept; two-sided pairwise *P* values were Holm-Bonferroni-adjusted across the six comparisons (wild type versus hypo-P, *P* = 0.13; wild type versus PP1-mut, *P* = 0.0013; wild type versus hyper-P-mut, *P* = 3.4 × 10^-20^; hypo-P versus PP1-mut, *P* = 9.2 × 10^-7^; hypo-P versus hyper-P-mut, *P* = 4.8 × 10^-29^; PP1-mut versus hyper-P-mut, *P* = 3.3 × 10^-12^).

Using the corelet assay, we quantified condensation as the percentage reduction in soluble cytoplasmic SspB-mScarlet3-pp150 IDR fluorescence between the pre-and post-activation frames, which estimates the fraction of the pp150 IDR incorporated into condensates (condensation efficiency; Fig. 4b). Wild-type pp150 IDR was efficiently recruited upon activation. In contrast, hyper-P-mut failed to form detectable condensates, showing that added negative charge strongly suppresses pp150 IDR phase separation (P = 3.4 × 10⁻²⁰; Fig. 4c). PP1-mut also showed reduced condensation (P = 0.0013), consistent with increased phosphorylation disfavouring condensation. Since PP1-mut alters only the two short PP1-docking motifs, whereas hyper-P-mut replaces 32 residues, the shared loss of condensation cannot be explained by the compositional change of the IDR, but is instead caused by its phosphorylation state. Conversely, the hypo-P mutant showed a modest increase in recruitment relative to wild-type, although this did not reach statistical significance (P = 0.13; Fig. 4c). This most likely reflects a limitation of the corelet assay. The seed imposes a high local valency that already drives wild-type condensation efficiently, so the assay detects suppression well but leaves little room to detect enhancement. Whether reduced phosphorylation promotes condensation therefore has to be tested without artificial clustering, which we do below in infected cells. These results identify phosphorylation as a tunable input controlling pp150 condensation, and suggest that the balance of CDK and PP1 activity could modulate the assembly and disassembly of pp150-rich tegument phases during viral assembly.

### Loss of pp150 phosphorylation causes ectopic tegument assembly and alters tegument ultrastructure in infection

To establish the relevance of pp150 condensate phosphoregulation in the full infection-context, we introduced the corresponding pp150 phospho-mutants into the HCMV strain TB40. All 32 minimal S/T-P CDK consensus sites within the pp150 IDR were mutated either to alanine (hypo-P) or to glutamate (hyper-P-mut), using the same residue set as in the corelet assays.

The hypo-P mutant showed a reproducible replication defect in multistep growth curves across three independent experiments (35.7-fold reduction at 16 dpi; Fig. 5a), indicating impaired production of viral progeny. Since reduced titers can result from impaired viral gene expression rather than from disturbed tegument assembly, we compared viral protein accumulation between wild-type and mutant infection. Over a 96 h time course, we found no differences in the expression of pp150 or of representative immediate-early, early and late proteins (IE1, IE2, UL84, pp65, UL99), in protein abundance or kinetics, between wild-type and mutant (Supplementary Fig. 4), arguing against a global gene-expression defect or impaired accumulation of major tegument proteins as an explanation for the observed growth defect.

**Fig. 5:**
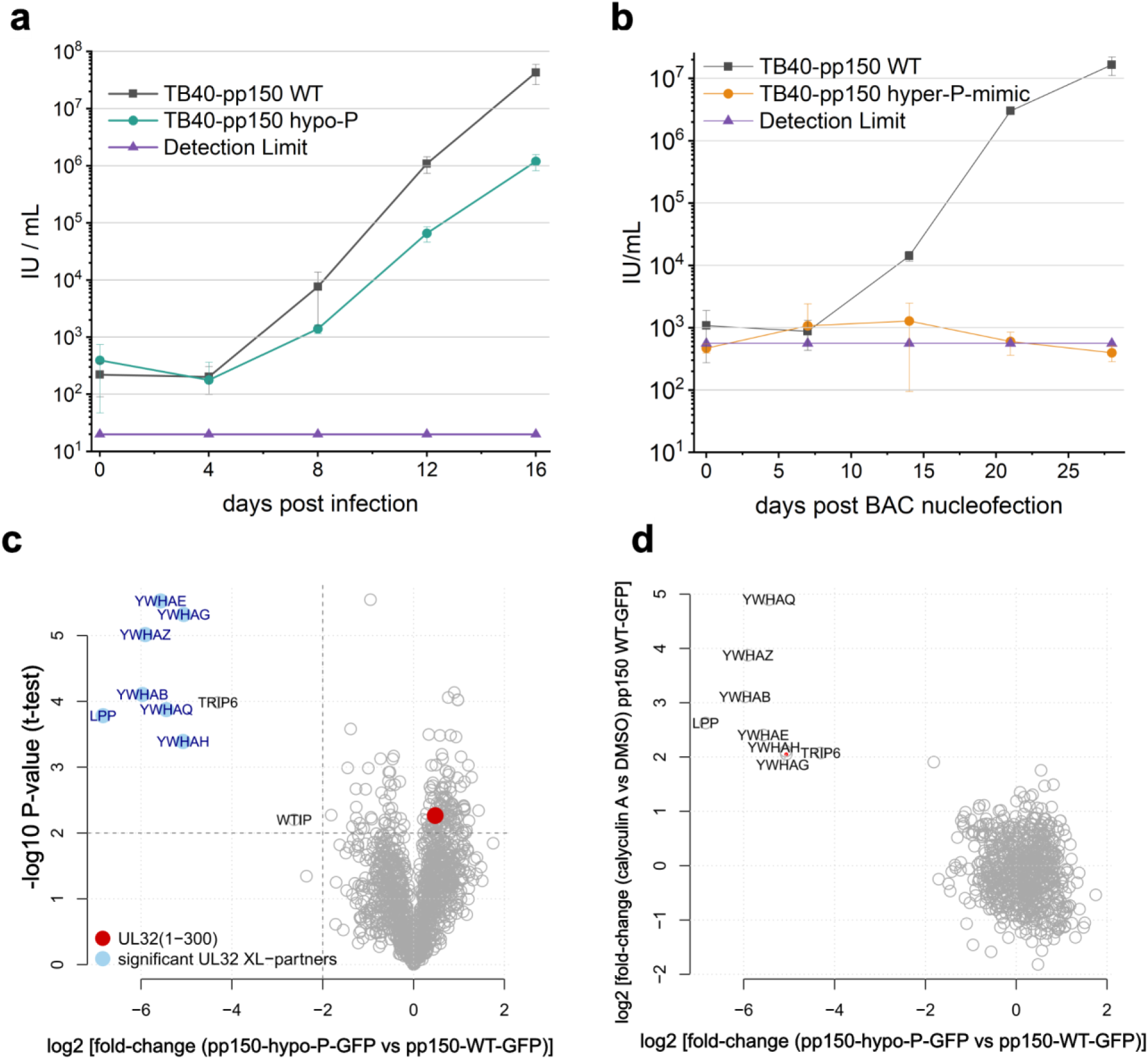
Hypophosphorylated pp150 impairs viral replication in a phosphorylation-dependent manner. **a**, Multistep replication kinetics of the hypo-P mutant (green) and wild-type (grey) virus in MRC-5 cells infected at a multiplicity of infection (MOI) of 0.01. Supernatant titers were determined at four-day intervals until day 16. Points, mean of three independent experiments (n = 3); error bars, standard deviation. Detection limit (19.8 IU/mL, magenta). **b**, The pp150 hyper-P mutant yielded no recoverable infectious progeny. Points, mean titer of three independent infections (n = 3); error bars, standard deviation. Detection limit (560 IU/mL, magenta). **c**, Volcano plot of affinity purification-mass spectrometry data comparing the pp150 interactome in hypo-P and wild-type infected cells from three independent infections (n = 3). x axis, log2 fold change of proteins enriched with hypo-P relative to wild-type pp150; y axis, -log10 P value (two-sided *t* test). **d**, Comparison of phosphorylation-dependent changes in the pp150 interactome. Log2 fold changes between hypo-P and wild-type infected cells are plotted against log2 fold changes reported in a published dataset for calyculin A treatment of wild-type infected cells relative to a vehicle control. The calyculin A dataset was not generated in this study. This analysis relates interactome changes caused by the hypo-P mutation to those induced by inhibition of PP1-and PP2A-dependent dephosphorylation.

We next asked whether these mutations affected the pp150 interactome, and in particular whether the hypo-P substitutions act through phosphorylation or through the compositional change they necessarily introduce. Label-free affinity purification-mass spectrometry (AP-MS) comparing the hypo-P mutant with wild type showed reduced association specifically with 14-3-3 and LIM-domain-containing proteins (Fig. 5c). 14-3-3 proteins are canonical phospho-serine/threonine readers, so their loss is the signature expected of reduced phosphorylation rather than of an arbitrary sequence change. We tested this directly by cross-comparison with a published interactome of wild-type pp150 under calyculin A treatment, a PP1/PP2A inhibitor^23^. The interactors lost in the hypo-P mutant were the same interactors that are sensitive to phosphatase inhibition (Fig. 5d). The hypo-P substitutions therefore act on phospho-dependent interactions, and the mutant is not characterized by a global remodelling of the pp150 interactome. Taken together, the hypo-P mutant has a defect in viral biogenesis that is not explained by a perturbed gene expression program or by non-specific disruption of pp150 interactions.

We next examined tegument organization directly using immunofluorescence microscopy. In hypo-P-infected cells, pp150 accumulated in large perinuclear assemblies, often on the opposite side of the nucleus from the viral assembly compartment (Fig. 6a). These aberrant pp150-positive assemblies were present in 90.8% of hypo-P-infected cells, compared with approximately 8.6% of wild-type-infected cells across three independent replicates (P = 3.8 × 10⁻⁴; Fig. 6b). Because the assemblies were also positive for 4′,6-diamidino-2-phenylindole (DAPI) (Fig. 6a), we labelled newly synthesized DNA with 5-bromo-2′-deoxyuridine (BrdU), which produced strong signals within the assemblies, indicating that they contain viral DNA (Supplementary Fig. 5).

**Fig. 6:**
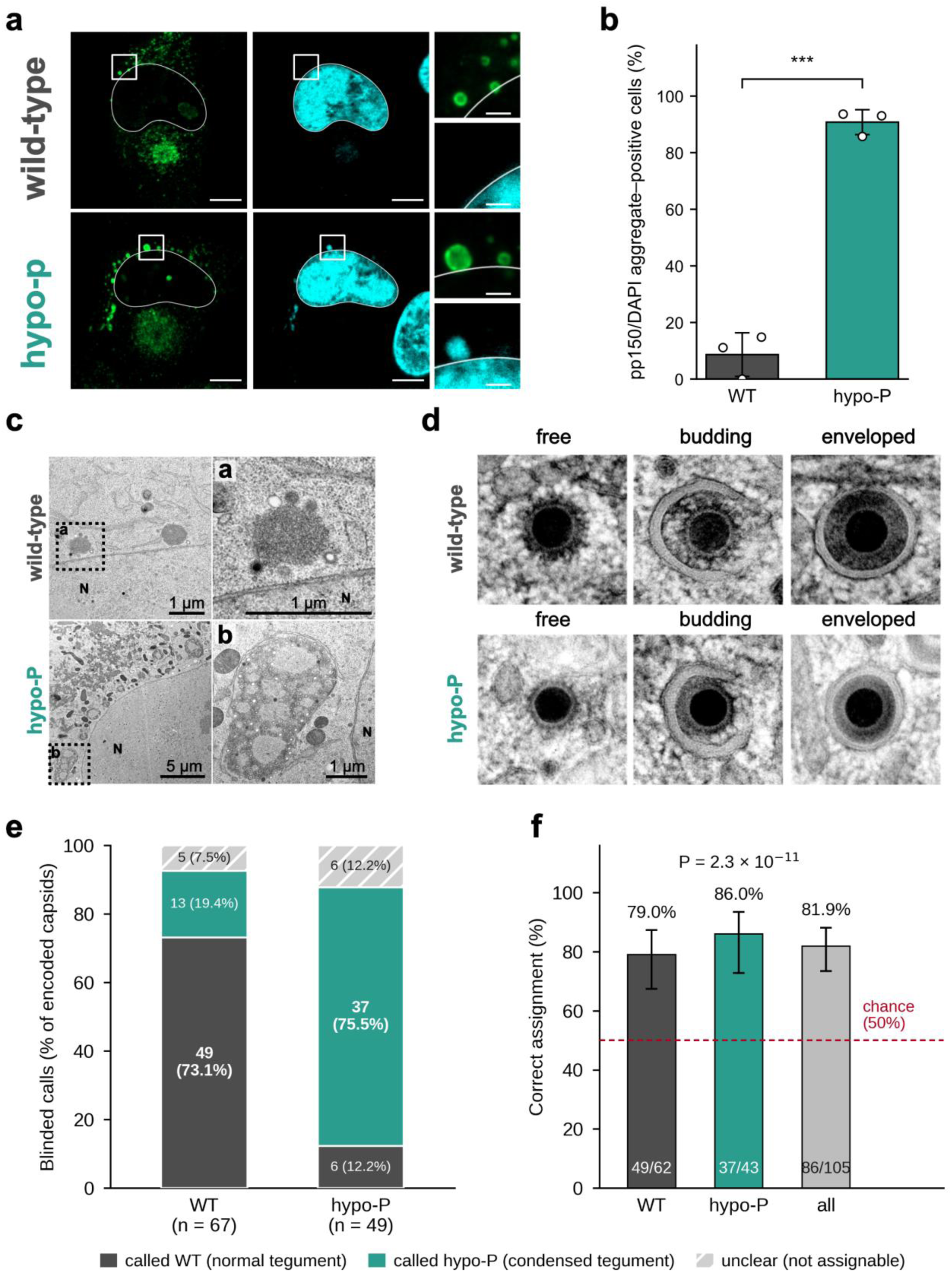
Hypophosphorylated pp150 drives aberrant cytoplasmic aggregation of tegument and capsids. **a**, Indirect immunofluorescence of human foreskin fibroblasts (HFFs) infected with the indicated viruses at an MOI of 1, fixed at 120 h post infection (hpi). pp150, green; nuclei stained with DAPI, blue, and outlined in white. Scale bars, 10 µm (overviews) and 2 µm (higher-magnification insets). **b**, Percentage of cells scored as positive or negative for DAPI-positive cytoplasmic aggregates. Bars, mean of three independent experiments (n = 3); error bars, standard deviation; *P* = 3.79 × 10^-4^, unpaired two-sided Welch t-test. **c**, Electron micrographs of wild-type (top) and hypo-P (bottom) infected HFFs at 120 hpi. Insets show capsid-associated protein aggregates close to the nuclear membrane. N, nucleus. **d**, Electron micrographs of individual capsids undergoing tegumentation and envelopment within the cytoplasmic viral assembly compartment. **e**, Distribution of blinded classifications of all coded capsids per true genotype. Segments, fraction of capsids called wild type (normal tegument), hypo-P (condensed tegument) or unclear (light grey, hatched); numbers within segments give capsid counts and percentages. n = 67 wild-type and 49 hypo-P capsids from one biological replicate. **f**, Fraction of classifiable capsids assigned to the correct genotype, for wild type, hypo-P and all capsids combined. Error bars, 95% Wilson score intervals; dashed line, 50% chance level; two-sided exact binomial test of the overall accuracy against chance, *P* = 2.3 × 10^-11^ (shown); true and blinded genotype were additionally associated by two-sided Fisher’s exact test, *P* = 3.1 × 10^-11^; Cohen’s κ = 0.64.

Transmission electron microscopy (TEM) revealed large cytoplasmic accumulations containing clusters of nucleocapsids embedded in an electron-dense matrix consistent with tegument material with membranous structures apparently wrapping around them, a phenotype absent from wild-type infection (Fig. 6c, Supplementary Fig. 6). Thus, phospho-ablation of the pp150 IDR produced aberrant perinuclear assemblies containing non-enveloped capsids and tegument material.

The residual capsids that underwent tegumentation and envelopment at their correct location within the viral assembly compartment displayed a subtly altered tegument, with a smoother and more condensed texture (Fig. 6d). To establish that this difference was reproducible rather than a product of observer expectation, individual capsids of both genotypes were assigned random codes and classified by an EM-experienced observer blinded to genotype as showing either normal (wild-type-like) or condensed (hypo-P-like) tegument, with an explicit “unclear” category for particles that could not be assigned. Among the 105 classifiable capsids, 86 (81.9%) were assigned to the correct genotype. Discrimination was successful in both directions: 49 of 62 wild-type capsids (79.0%) and 37 of 43 hypo-P capsids (86.0%) were correctly identified (P = 2.3 × 10⁻¹¹ against chance; Cohen’s κ = 0.64; Fig. 6e, f).

Importantly, the hyper-P mutant yielded no detectable infectious progeny at any time point up to 28 days post BAC nucleofection (Fig. 5b). Because the phosphomimetic IDR did not condense even at capsid-like density in cells (Fig. 4b, c), it is likely that these virions cannot nucleate a tegument condensate around the capsid, as predicted by a model in which condensation is required for tegument assembly.

Together, these results show that phospho-specific manipulation of the pp150 IDR alters tegument behavior in infection as predicted by our corelet condensation assays, and it does so in both directions. Insufficient phosphorylation drives aberrant cytoplasmic condensation of tegument material around capsids, consistent with a loss of phosphorylation-dependent control over pp150, whereas constitutive phosphorylation, which abolished condensation in cells, is incompatible with the production of infectious progeny.

## Discussion

HCMV tegument assembly and final envelopment are well described morphologically, but their molecular coordination remains poorly understood. A central question is how an amorphous protein layer selectively assembles around capsids and engages the membranes required for envelopment. Our findings identify pp150 as a phosphorylation-regulated condensate scaffold that recruits additional tegument proteins and associates with membrane-bound partners, providing a mechanistic framework for coordinating these steps.

Here, we show that the IDR of pp150 forms liquid-like condensates in living cells when locally clustered, mimicking the high local concentration of pp150 on the capsid surface. Many studies of LLPS rely on purified proteins in vitro, which provides biochemical control but does not fully capture the intracellular environment. In contrast, the corelet system allowed us to test pp150 phase behavior in a cellular context, where molecular crowding, ion composition, membranes, and potential cellular interaction partners are present and can influence condensate formation. Although this heterologous system does not recapitulate HCMV infection, it demonstrates that the pp150 IDR has an intrinsic capacity for condensation when concentrated above a threshold.

This finding fits with an expanding body of work showing that LLPS contributes to diverse cellular processes and is used by viruses to organize replication and assembly. In herpesviruses, phase separation has been implicated in the behavior of herpes simplex virus 1 (HSV-1) tegument proteins and in the formation of HCMV replication compartments and γ-herpesvirus assembly compartments^19–21^, but none of these studies could link condensate behavior in cells to tegument assembly in infection and show that a tegument protein forms a phosphorylation-tunable condensate which recruits other tegument proteins and engages membranes, and that disrupting this behavior in recombinant virus misplaces tegument assembly and blocks virion production.

For HCMV tegument assembly, pp150 condensation provides a mechanistic framework for how a protein layer can form around capsids. Because pp150 is anchored to the capsid through its structured N-terminal region^22^, its large C-terminal IDR could generate a capsid-associated condensate that supports multivalent and robust recruitment of additional tegument proteins, as we show here for UL71 and UL25. This model is further strengthened by our recent finding that the pp150 IDR appears as a major interaction hub within the HCMV tegument in a crosslinking-based study^23^.

A major finding of this work is that pp150 condensates can interact with membrane-associated UL71, thereby linking capsid-bound tegument to the membranes used for envelopment. This not only establishes a direct capsid-to-membrane axis but also introduces a conceptual framework for addressing open questions about the final envelopment of HCMV. Recent work has shown that condensates can interact with lipid membranes through wetting, local membrane remodeling, and, in some systems, membrane bending, engulfment, and scission-like processes, mechanisms that resemble the membrane events of HCMV envelopment^15–18^. The association of a tegument-derived condensate with UL71-positive membranes gives a defined point of contact between tegument and membrane, and makes it testable whether condensate wetting helps to bend, wrap, and sever these membranes around the capsid.

Furthermore, our results show that pp150 phase separation is regulated by phosphorylation. More broadly, phosphorylation is a well-established regulator of LLPS, particularly for IDR-containing proteins, where changes in charge, interaction valency, and binding-partner recruitment can strongly alter condensate formation and condensate properties^13,14^. A phosphorylation-based regulatory model is also consistent with previous observations during HCMV infection. PP1 is upregulated during infection and incorporated into the viral particle, suggesting that local dephosphorylation may contribute to late-stage tegument maturation^31,32^. Conversely, inhibition of specific kinases has been reported to induce cytoplasmic accumulations of tegument proteins^33^.

Such regulation is likely essential during assembly, because pp150 condensation must occur at the correct time and place, consistent with emerging evidence for progressive tegument formation during final envelopment^34^. This is supported by the phenotype of the hypo-P mutant, in which pp150 accumulated in aberrant cytoplasmic assemblies that contained viral DNA and capsids. Insufficient phosphorylation thus appears to release pp150 condensation from its normal spatiotemporal control, leading to uncontrolled condensation, plausibly through increased self-association of the pp150 IDR.

Together, our findings support a model in which pp150 acts as a regulated condensate scaffold that organizes the HCMV tegument and couples it to membranes during final envelopment. This suggests that condensate-based assembly is not unique to HCMV, but represents a broader strategy for building tegument layers from multivalent, IDR-rich proteins in the absence of extensive long-range structural order. Indeed, in an accompanying manuscript, we identify the herpes simplex virus 1 (HSV-1) protein VP22 as a condensate-forming scaffold that organizes the HSV-1 tegument through a comparable mechanism^26^. VP22 shares no detectable sequence relationship with pp150 and, unlike pp150, does not bind capsids directly, yet it performs an equivalent organizing function. Given that the alphaherpesvirus lineage separated from the lineage leading to the beta-and gammaherpesviruses an estimated 180 to 220 million years ago^35^, this shared behavior is unlikely to reflect conservation of a single protein, and instead suggests that convergent evolution has settled on the same physical mechanism for tegument formation. Possibly, condensation provides a robust avenue to concentrate a compositionally variable protein layer around the capsid while allowing metastability for subsequent disassembly, potentially via phosphoregulation.

## Materials and Methods

### Cells and culture conditions

VeroB4 cells (DSMZ, ACC 33) were used for all transfection experiments and were maintained in Dulbecco’s modified Eagle medium (DMEM; Gibco) supplemented with 10% (v/v) fetal calf serum (FCS; Gibco). Human foreskin fibroblasts (HFFs; isolated from anonymous voluntary donors with their informed consent) and human embryonic lung fibroblasts (MRC-5; ATCC CCL-171) were used for virus propagation and infection experiments. MRC-5 cells were maintained in Eagle’s minimum essential medium (EMEM; Merck M2279) supplemented with Earle’s balanced salt solution, 25 mM HEPES, 1 mM sodium pyruvate, 2 mM L-alanyl-L-glutamine, non-essential amino acids (NEAA), 0.75‰ (w/v) sodium bicarbonate, 50 µg/mL gentamicin, and 10% (v/v) fetal bovine serum (FCS). HFFs were maintained in fibroblast culture medium consisting of DMEM (Thermo Fisher Scientific) supplemented with 100 units/mL penicillin, 100 µg/mL streptomycin, 10% (v/v) FCS, and 1× NEAA. All cell lines were cultivated at 37 °C and 5% CO₂ in a humidified incubator and were routinely tested negative for mycoplasma contamination (MycoAlert Mycoplasma Detection Kit, Lonza).

### Viruses and infection conditions

Variants derived from the human cytomegalovirus (HCMV) strain TB40-BAC4 were used for all infection experiments^36^. A detailed list of all viruses used is provided in Supplementary Table 1a. Infectious titers of cell-free virus stocks were determined by immunotitration and are expressed as immediate-early-forming units (IU)/mL, as previously described^37^. Infections were performed at a multiplicity of infection as indicated for each experiment. Before infection, fibroblasts were synchronized in the G0/G1 phase of the cell cycle by omitting FBS from the culture medium for up to 48 h.

### Plasmid cloning and transfection

Plasmids were generated by In-Fusion cloning (Takara), and all constructs were sequence-verified. Detailed information for each construct is provided in Supplementary Table 1b.

VeroB4 cells were seeded into µ-Slide 8-well chamber slides (ibidi, 80806) one day before transfection to reach approximately 60-70% confluency at the time of transfection. Transfections were performed with Lipofectamine 3000 (Thermo Fisher Scientific) according to the manufacturer’s instructions with minor adjustments. Per well, 360 ng total plasmid DNA and 1 µL P3000 reagent were diluted in 15 µL Opti-MEM. In parallel, 0.9 µL Lipofectamine 3000 was diluted in 15 µL Opti-MEM. The two solutions were combined, mixed gently, incubated for 10 min at room temperature, and added dropwise to each well. Plasmid ratios were held constant across experiments. Where two plasmids were co-transfected, they were combined at a 1:1 ratio (180 ng each). Where the corelet components iLID-eGFP-FTH1 and SspB-mScarlet3-pp150 IDR were co-transfected with UL71-SNAP or UL25-SNAP, the three plasmids were combined at a 2.5:2.5:1 ratio (150 ng, 150 ng, and 60 ng, respectively). Cells were imaged at 24 hours post transfection.

### BAC mutagenesis and virus reconstitution

The bacterial artificial chromosome (BAC)-derived hypo-P mutant was generated by two-step Red-mediated mutagenesis as described^38^. DNA sequences used for mutagenesis are listed in Supplementary Table 1c. To generate the hypo-P mutant virus, carrying 32 serine/threonine-to-alanine substitutions within the C-terminal intrinsically disordered region (IDR) of pp150, we first constructed a transfer plasmid enabling en bloc insertion of the mutated IDR into the viral genome. For this purpose, the Kanamycin-SceI (KanS) cassette from pEP-KanS2^39^ was PCR-amplified using primers UL32_Kn_in_dupl_SAC2+ and UL32_Kn_in_SAC2-and inserted into pET52b(+)-pp150c^24^ via SacII restriction sites, yielding pET52b(+)-pp150c-KanS. In this plasmid, the pp150 wild-type sequences flanking the KanS cassette were replaced with synthetic mutated pp150 fragments (gBlocks, Integrated DNA Technologies). Fragment pp150-SPTPless-partA, encompassing pp150 codons 303-702 and containing four SP→AP and seven TP→AP substitutions, was cloned into the XbaI/XhoI sites of pET52b(+)-pp150c-KanS. Fragment pp150-SPTPless-partB, spanning pp150 codons 685-1049 and carrying twelve SP→AP and eight TP→AP substitutions, was inserted via HindIII/SacI restriction sites. Similarly, for generation of the hyper-P mimic pp150 mutant, synthetic fragments pp150-S/TPtoEP-partA and pp150-S/TPtoEP-partB were cloned into XbaI/XhoI and HindIII/SacI sites respectively. A linear PCR fragment containing the complete mutated pp150-IDR (codons 303-1049) together with the KanS cassette was subsequently amplified using primers UL32Cmut+Kana-recomb-fw and UL32Cmut+Kana-recomb-rv for recombination into TB40-BAC4. To facilitate efficient and seamless integration of the mutated pp150 region, the corresponding wild-type sequence in TB40-BAC4 encompassing pp150 codons 303-1049 was first deleted by traceless BAC mutagenesis, using primers UL32(303-1049)-del-fw and UL32(303-1049)-del-rv for KanS amplification and genomic insertion. To introduce an additional SP→AP substitution at codon 294 of the hypo-P mutant, primers UL32-S294A-fw and UL32-S294A-rv were used. The transfer plasmid pEP-EGFPin (gift from Nikolaus Osterrieder, Addgene plasmid #60961; http://n2t.net/addgene:60961; RRID:Addgene_60961) was used for the insertion of C-terminal EGFP tags into pp150 using the PCR primers UL32-Cterm-EGFP_fw and UL32-3UTR-EGFP_rv. To generate monomeric EGFP (mEGFP), the A206K mutation was introduced into the EGFP sequence using the primers EGFP-A206K-fw and EGFP-A206K-rv. All mutations were verified by PCR and Sanger sequencing. In addition to the intended substitutions, the hypo-P pp150 gene carried an unintended Thr-to-Ala point mutation at amino acid position 932. Recombinant virus was reconstituted by co-transfecting purified 3.3 µg BAC-DNA and 0.7 µg pp71 expression plasmid into 1.0E+06 MRC-5 cells by nucleofection with Nucleofector^TM^ II (amaxa biosystems) and in-house made buffer^40^.

### Live-cell imaging and optogenetic corelet activation

Live-cell fluorescence imaging was performed on two spinning-disk confocal systems of similar but not identical configuration. Both were built around a Nikon Eclipse Ti2 inverted microscope equipped with a Yokogawa CSU-W1 spinning-disk unit and were operated under NIS-Elements AR. System 1 was equipped with an Andor iXon Ultra DU-888U3 EMCCD camera, a 100×/1.45 NA oil-immersion objective, and a Fluorescence Recovery after Photobleaching (FRAP) unit from Rapp Optoelectronics. System 2 was equipped with two Hamamatsu ORCA-Fusion BT sCMOS cameras (C14440-20UP) and an Apo TIRF 100×/1.49 NA oil-immersion objective. On both systems, cells were maintained at 37 °C and 5% CO₂ using a stage-top incubator.

System 1 was used for the FRAP experiments and for the quantitative measurement of corelet condensation efficiency. System 2 was used for all remaining live-cell experiments, including the qualitative condensate-formation screen and the co-condensation experiments with UL25-SNAP, UL71-SNAP, and UL99-SNAP.

Corelet activation was induced by 488 nm illumination, which simultaneously excited the iLID-eGFP-FTH1 seed construct and triggered the light-dependent iLID-SspB interaction. No separate activation pulse was therefore applied, and activation occurred during image acquisition of the eGFP channel. To ensure that the first time point contained pre-activation information for the remaining fluorescent channels, the 488 nm/eGFP channel was acquired last in each imaging cycle. This acquisition order recorded the SspB-mScarlet3-pp150 IDR and SNAP-tagged constructs before blue-light activation at the first frame, while subsequent time points captured the dynamics after activation.

SNAP-tagged constructs were labeled before imaging using SNAP-Cell® 647-SiR or SNAP-Cell® 505-STAR substrates (New England Biolabs) according to the manufacturer’s protocol. Briefly, cells were incubated with the respective SNAP substrate at 37°C, washed with pre-warmed media to remove unbound fluorophore and imaged immediately thereafter. For membrane labeling, FM4-64FX (Invitrogen, F34653) was added directly to the imaging medium (final concentration 5 μg/mL) and cells were imaged immediately after dye addition.

### Quantification of corelet condensation efficiency

Condensation of pp150 IDR variants was quantified in the optogenetic corelet system from time-lapse movies acquired under identical imaging conditions in transfected VeroB4 cells. For each cell, three circular cytoplasmic regions of interest (ROIs) that were devoid of condensates in the final post-activation frame were drawn manually, using the visible contrast between diffuse cytoplasmic background and condensate foreground as the selection criterion. Mean mScarlet3 fluorescence intensities within these ROIs were measured in the first (pre-activation) and last (post-activation) frame. Background fluorescence, determined from a signal-free area, was subtracted from all measurements, and intensities were normalized to the mean fluorescence of a larger cytoplasmic reference region to correct for photobleaching. The resulting decrease in soluble cytoplasmic mScarlet3 signal was interpreted as incorporation of SspB-mScarlet3-pp150 IDR into surrounding condensates and is reported per cell as the percentage reduction in cytoplasmic fluorescence. Measurements were made in NIS-Elements AR Analysis software (version 6.10.02, Nikon).

The hyper-P-mut variant did not produce visible condensates under any condition tested. For this construct, evenly fluorescent cytoplasmic regions were therefore selected, so that the resulting values report the noise floor of the measurement rather than incorporation of pp150 IDR into condensates, and fewer cells were sampled accordingly. The experimental unit was the individual cell, nested within the biological replicate. For each construct, cells were analyzed across three independent biological replicates: wild-type, hypo-P, and PP1-mut, 20 cells per replicate, n = 60 cells; hyper-P-mut, 10 cells per replicate, n = 30 cells.

Condensation values were analyzed with a linear mixed-effects model, because individual cells are nested within biological replicates and are therefore not statistically independent. The three cytoplasmic ROIs measured per cell were averaged before analysis, so that each cell contributed a single value. The model included construct (wild-type, hypo-P, PP1-mut, hyper-P-mut) as a fixed effect and biological replicate as a random intercept, with wild-type as the reference level, and each cell contributed one measurement. Models were fitted by restricted maximum likelihood (REML) using statsmodels (version 0.14, Python 3.12.3). The overall effect of construct was assessed with a joint Wald test of the null hypothesis that all construct coefficients equal zero. Pairwise differences were then estimated by refitting the same mixed-effects model to each pair of constructs separately. The resulting P values were adjusted across the six pairs by the Holm-Bonferroni procedure to control the family-wise error rate. Variance components were inspected to confirm that the random-effect structure was appropriate. Between-replicate variance was negligible relative to cell-to-cell variance (intraclass correlation coefficient, ICC, approximately 0), indicating that biological replicates were highly reproducible.

### Fluorescence recovery after photobleaching

Fluorescence recovery after photobleaching (FRAP) experiments were performed on the spinning-disk confocal system described above. Individual condensates were selected and photobleached using a 473 nm laser. Fluorescence recovery within the bleached region was monitored for 60 s at 5 frames per second under live-cell conditions. Quantitative analysis was performed using NIS-Elements AR Analysis software (version 6.10.02, Nikon). Mean fluorescence intensity within the bleached region of interest (ROI) was measured over time. Background fluorescence was subtracted, and intensity values were normalized to an internal, unbleached reference region within the same cell to correct for acquisition-induced photobleaching. Pre-bleach intensity was set to 100%. For each biological replicate (n = 3), 10 cells were analyzed and averaged.

Recovery curves were fitted to a nonlinear regression model (ExpAssoc1) in OriginPro (version 2025b, OriginLab) and used to calculate the half-time of recovery and the mobile fraction. The recovery plateau was calculated as F∞ = Yb + A, where Yb is the fluorescence intensity immediately after bleaching and A is the amplitude of recovery from the nonlinear fit. The mobile fraction was determined as (F∞ - Yb) / (100 - Yb). Values are reported as mean ± standard deviation of three biological replicates.

### Disorder-enrichment analysis of pp150 contacts

Crosslinks involving pp150 were extracted from a published crosslinking mass spectrometry (XL-MS) dataset of HCMV virions and mapped onto the protein sequences of HCMV strain TB40-E clone BAC4 (GenBank accession EF999921)^23^. The UL45 sequence, which is not annotated in the EF999921 record, was obtained from the TB40-BAC4^36^ sequence and verified against the crosslink coordinates. Per-residue intrinsic disorder was predicted using IUPred3 in long mode, run locally^29^. For each partner protein, an interface region was defined as a window of ±7 residues centered on each lysine crosslinked to pp150. Envelope glycoproteins were included among the tegument-facing proteins, because their pp150-crosslinked residues face the tegument compartment inside the viral membrane. Capsid and capsid-associated proteins (UL46, UL48A, UL77, UL80, UL85, UL86, UL93) were analyzed as a separate contrast group, because they are structurally distinct from the tegument. Host proteins crosslinked to pp150 were analyzed as a second contrast group.

The mean disorder of the pp150-contact windows was compared with that of background windows centered on the remaining crosslinked lysines of the same protein, those are, lysines crosslinking to partners other than pp150 in the full virion crosslink dataset. Because all background lysines are themselves observed in crosslinks and are therefore solvent-exposed and reaction-competent, this accessibility-matched, within-protein design controls for crosslinker lysine reactivity, solvent accessibility and distance constraints, protein abundance, and protein length. Proteins with at least one pp150-contacting lysine and at least two other crosslinked lysines were included in the test. Proteins lacking a sufficient within-protein background were excluded from the significance testing. Δdisorder was defined as contact-window disorder minus background-window disorder, such that positive values indicate greater disorder at pp150 contact sites than at background crosslinked lysines of the same protein.

The experimental unit was the protein. Individual lysine-centered windows were averaged within each protein before testing. Normality of the paired differences was assessed by Shapiro-Wilk. Because normality was rejected for the tegument class (Shapiro-Wilk *W* = 0.866, *P* = 0.0295), the two-sided Wilcoxon signed-rank test on the within-protein difference was used as the primary test for all classes. A one-sample t-test against zero was computed on the same values as a sensitivity analysis and is reported in the Source Data file. In no case did the parametric and non-parametric tests lead to different conclusions. Δdisorder was compared between the tegument-facing, capsid and host classes by Mann-Whitney U test. These between-class tests were one-sided, reflecting the directional hypothesis, specified before analysis, that disorder is enriched at pp150 contact sites in tegument-facing proteins relative to capsid and host proteins.

### Multistep growth kinetics

Viral growth of the pp150 hypo-P mutant was assessed in a low multiplicity of infection (MOI) multistep growth analysis with three independent replicates. MRC-5 cells were infected with wild-type or hypo-P virus at an MOI of 0.01. Supernatants were collected at four-day intervals until day 16 and were then used to infect MRC-5 cells. The number of infected cells from these infections was determined at 24 h post infection by flow-cytometric quantification of immediate-early (IE) protein-expressing cells, as previously described^37^. The detection limit was determined by measuring the background of falsely detected infected cells in a sample of uninfected cells.

Because no infectious virus stock of the pp150 hyper-P mimic mutant could be recovered, its replication could not be assessed under MOI-controlled conditions. Growth analysis of this mutant was therefore initiated directly from BAC nucleofection, performed as described above, comparing wild-type and hyper-P-mut BAC in parallel. Supernatants were collected at seven-day intervals until day 28 post nucleofection and titrated on MRC-5 cells as described above, using flow-cytometric quantification of IE protein-expressing cells at 24 h post infection and the same determination of the detection limit. Three independent nucleofection experiments were performed.

### Immunoblot analysis

MRC-5 cells were infected at an MOI of 5 IU/cell and harvested at the indicated time points by trypsinization. Cell pellets were lysed by sonication in lysis buffer containing 50 mM Tris-Cl (pH 6.8), 2% sodium dodecyl sulfate (SDS), 10% glycerol, 1 mM dithiothreitol (DTT), 2 µg/mL aprotinin, 10 µg/mL leupeptin, 1 µM pepstatin, and 0.1 mM Pefabloc. Lysates were clarified by centrifugation, and protein concentrations were normalized using the Bio-Rad DC protein assay. Bromophenol blue and DTT (100 mM final concentration) were added before boiling the samples at 95 °C for 3 min. Proteins were separated by SDS-PAGE and transferred onto polyvinylidene fluoride (PVDF) membranes. Membranes were blocked in Tris-buffered saline containing 0.1% Tween-20 and 5% (w/v) skim milk to reduce non-specific binding.

The following primary antibodies were used: anti-IE1/IE2 (clone 8B1.2, Merck MAB810X, 1:2500), anti-UL84 (clone Mab84, Santa Cruz sc-56977, 1:500), anti-pp65 (clone 3A12, Santa Cruz sc-56973, 1:500), anti-UL99 (clone CH19, Santa Cruz sc-69749, 1:500), anti-pp150 (clone XP1, a gift from Bodo Plachter, 1:3000), and anti-RPS6 (clone 5G10, Cell Signaling 2217, 1:1000). Immunoblots were developed using horseradish peroxidase-conjugated secondary antibodies against mouse IgG (Agilent P044701-2, 1:2000) and rabbit IgG (Agilent P044801-2, 1:2000) in combination with enhanced chemiluminescence detection reagents (SuperSignal West Dura, Thermo Fisher Scientific 34075).

### GFP affinity purification

MRC-5 cells were infected with HCMV-pp150-mEGFP (WT) or HCMV-pp150-hypo-P-mEGFP at an MOI of 5 IU/cell. Experiments were performed in n = 3 replicates, with one confluent 15 cm dish as starting material per replicate and experiment. At 4 days post infection, cells were collected by scraping in PBS and processed as previously described^41^. Briefly, cells were washed in PBS and lysed for 20 min in lysis buffer (25 mM Tris-HCl (pH 7.4), 125 mM NaCl, 1 mM MgCl₂, 1% Nonidet P-40, 0.1% SDS, 5% glycerol, 1 mM dithiothreitol, 2 µg/mL aprotinin, 10 µg/mL leupeptin, 1 µM pepstatin, 0.1 mM Pefabloc, 0.5 mM Na₃VO₄, 10 mM β-glycerophosphate, 1 mM NaF). Lysates were sonicated to solubilize nucleocapsid-associated pp150-mEGFP before clearing for 20 min at 12,000 g at 4 °C.

For GFP affinity purification, µMACS anti-GFP microbeads (130-091-125, Miltenyi Biotec) were used. After loading onto µ-Columns (Miltenyi Biotec), lysis buffer was used for the first washing step, lysis buffer without detergent for the second, and 25 mM Tris-HCl (pH 7.4) for the final washing step. Proteins were eluted by adding 200 µL of 8 M guanidine hydrochloride pre-warmed to 95 °C. Proteins were precipitated from the eluates by adding 1.8 mL LiChrosolv ethanol (Merck) and 1 µL GlycoBlue (Thermo Fisher Scientific). After incubation at 4 °C overnight, samples were centrifuged for 1 h at 4 °C, ethanol was decanted, and the pellet was air-dried. Proteins were then resolved in digestion buffer (50 mM triethylammonium bicarbonate, pH 8.0, 1% sodium deoxycholate, 5mM tris(2-carboxyethyl)phosphine hydrochloride (TCEP) and 30 mM chloroacetamide (CAA)) and supplemented with trypsin and LysC at 1:25 and 1:100 enzyme-to-protein ratios (w/w), respectively. Digests were incubated overnight at 37 °C, subjected to C18 stage-tip desalting, and analyzed by liquid chromatography-mass spectrometry (LC-MS).

### LC-MS measurement of bottom-up proteomic samples

Bottom-up proteomic samples were analyzed on an Orbitrap Exploris 480 mass spectrometer (Thermo Fisher Scientific) coupled online to a Vanquish Neo UHPLC system (Thermo Fisher Scientific) and operated with instrument control software version 4.2. Peptides were loaded onto a 50 cm in-house packed reverse-phase analytical column (Poroshell 120 EC-C18, 2.7 µm, Agilent Technologies) and separated using a 120 min linear gradient of 0.1% formic acid in water (buffer A) and 0.1% formic acid in 80% acetonitrile (buffer B). MS1 spectra were acquired in the Orbitrap at a resolution of 120,000 with a cycle time of 2 s and a dynamic exclusion duration of 40 s. The precursor intensity threshold was set to 1E+4, the automatic gain control (AGC) target to 300%, and the maximum injection time to “Auto”. Precursors with charge states from +2 to +4 were isolated using a 1.6 m/z isolation window and fragmented by higher-energy collisional dissociation at a normalized collision energy of 30%. MS2 spectra were acquired in the Orbitrap at a resolution of 15,000 with the AGC target set to “Standard”.

### LC-MS data analysis

Raw DDA LC–MS/MS data were searched using the MSFragger search engine (v4.4.1) in the FragPipe version 24.0 environment^42^. Settings were set corresponding to the standard LFQ-phospho workflow, except that PTMProphet site localization was disabled and a minimum of two peptide ions was required for IonQuant protein quantification. Spectra were matched against in silico spectra from a human Uniprot reference proteome (downloaded 2020) combined with viral sequences of strain TB40/E clone BAC4 (GenBank: EF999921.1), common contaminants, the phosphosite-substituted sequence of pp150 and corresponding decoys with false discovery rate (FDR) set to 1% at the peptide-spectrum match (PSM), peptide, peptide-ion, and protein levels. For analysis, LFQ intensities were log2 transformed and Log2 fold-changes and p-values from a two-sided *t* test were calculated based on these values. The log2 fold-changes were subsequently compared to a previously published differential interactome of wild-type pp150 comparing treatment with the PP1/PP2A inhibitor calyculin A to DMSO-treated cells^23^.

### BrdU labeling and immunofluorescence

Nucleocapsids were detected by indirect immunofluorescence following pulse labeling of newly synthesized viral DNA with the thymidine analog 5-bromo-2′-deoxyuridine (BrdU, Thermo Fisher Scientific), as described^43^. HFFs seeded in µ-Slide 8-well chamber slides (ibidi) were infected with HCMV TB40 WT and HCMV-pp150-hypo-P at an MOI of 1. At 96 h post infection (hpi), infected cells were incubated with 10 µM BrdU in fibroblast culture medium at 37 °C for 20 h (pulse), before the BrdU-containing medium was replaced with fresh serum-reduced fibroblast culture medium (1% FCS) for 4 h (chase). At 120 hpi, cells were washed once with PBS and fixed with 4% PFA for 10 min at 4 °C.

Indirect immunofluorescence staining was performed as described^43^. Cells were permeabilized with 1% Triton X-100 (BrdU-labeled samples) or 0.1% Triton X-100 (samples without BrdU labeling) in PBS for 10 min at room temperature, then blocked with a solution containing 1% (w/v) BSA and 10% (v/v) horse serum in PBS for 30 min at room temperature. Primary and secondary antibodies were diluted in blocking solution. Cells were incubated overnight at 4 °C with the primary antibody, washed three times with washing solution (1% BSA and 0.1% Triton X-100 in PBS), and incubated with the secondary antibody for 45 min at room temperature. Cell nuclei were stained with 0.33 µg/mL 4′,6-diamidino-2-phenylindole (DAPI, Merck KGaA).

Cellular and viral protein stainings were performed before the detection of BrdU. Stained cells were incubated with 4% PFA for 15 min at 4 °C to preserve the staining against the DNA denaturation required for BrdU detection. Cells were then incubated with 2 M HCl for 15 min at room temperature to expose the incorporated BrdU residues for antibody staining, washed three times with PBS, and blocked with blocking solution for 30 min at room temperature. Cells were subsequently incubated with the primary antibody against BrdU and with the secondary antibody as described above.

The following antibodies were used. The HCMV cytoplasmic viral assembly compartment (cVAC) was detected with a mouse monoclonal antibody (MAb) against the cis-Golgi protein GM130 (clone 35/GM130, IgG1, BD Biosciences, 610823, 1:1000). pp150 was detected with a mouse antibody against pp150 (clone 36-14, IgG2b; kindly provided by William Britt, University of Alabama at Birmingham, USA, 1:1000). BrdU-labeled viral genomes were detected with a rat anti-BrdU MAb (clone RF06, Bio-Rad Laboratories, MCA6144, 1:500).

Representative cells from at least two independent experiments were selected for confocal microscopy. Confocal images were acquired with a 63× objective on an Axio Observer.Z1 fluorescence microscope equipped with an ApoTome 2.0 (Zeiss). Images were processed with ZEN 3.0 (blue edition) software (Zeiss).

### Quantification of aberrant pp150 aggregates

Human foreskin fibroblasts (HFF) were seeded in 18-well µ-slides (ibidi) and grown to confluence. Cells were serum-starved in medium containing 1% fetal bovine serum (FBS) for 24 h prior to infection and were then infected with approximately 50 immediate early (IE) units per well of either wild-type or hypo-P virus. Three independent infections were performed per virus. At 5 days post infection, cells were fixed, permeabilized, and stained with antibodies directed against the tegument protein pp150 and against the Golgi matrix protein GM130, as described above. Nucleic acids were counterstained with 4′,6-diamidino-2-phenylindole (DAPI).

Images were acquired with a 20× objective on an Axio Observer.Z1 fluorescence microscope (Zeiss) equipped with an ApoTome 2.0 (Zeiss) and analyzed in ZEN 3.0 (blue edition, Zeiss). Late-stage infected cells were identified by a sufficient cytoplasmic pp150 signal and were scored as aggregate positive or aggregate negative. Cells in which cytoplasmic pp150 colocalized with the GM130 signal, that is, was confined to the cytoplasmic viral assembly complex (cVAC), were scored as negative, whereas cells showing pp150 in discrete perinuclear aggregates outside the cVAC were scored as positive. Scoring was performed on randomly selected fields, and a total of 74 (wild type) and 139 (hypo-P) infected cells were scored across the three replicates (20 to 49 cells per replicate and condition).

Statistical analysis used the biological replicate, not the individual cell, as the experimental unit (n = 3 per condition). The percentage of aggregate positive cells per replicate was compared between conditions by an unpaired two-sided Welch t test.

### Transmission electron microscopy

Samples for transmission electron microscopy (TEM) were prepared by high-pressure freezing (HPF), freeze substitution, and Epon embedding, as previously described^43^. HFFs were seeded in µ-Slide 8-well chamber slides (ibidi) containing carbon-coated sapphire disks (3 mm diameter, 50 µm thickness; Engineering Office M. Wohlwend) and infected with wild-type or hypo-P virus at an MOI of 1. At 120 hpi, infected cells on sapphire disks were immobilized by HPF using a Compact 01 high-pressure freezer (Engineering Office M. Wohlwend). Samples were subsequently freeze-substituted and embedded in Epon. Ultrathin sections of 70 nm were cut from the Epon block with an EM UC7 ultramicrotome (Leica Microsystems) equipped with a 45° diamond knife and placed on Formvar-coated single-slot copper grids (Plano). Thin sections were examined with a JEM-1400 transmission electron microscope (JEOL) equipped with a Veleta charge-coupled device camera (Olympus) and operated at an acceleration voltage of 120 kV. The remaining infected cells in the µ-Slide 8-well chamber slides were fixed with 4% PFA and subjected to indirect immunofluorescence staining to control for infection rate and virus mutant phenotypes.

### Blinded classification of capsid-associated tegument

To assess whether the hypo-P mutant displays an altered capsid-associated tegument, HFF were infected with wild-type or hypo-P virus and processed for TEM as described above. At 5 days post infection, individual capsids at various stages of tegumentation and envelopment, including fully enveloped particles, located within the viral assembly compartment were selected at random and imaged at 80,000× nominal magnification. In total, 116 single capsid images were acquired (67 wild-type, 49 hypo-P) from 10 cells per genotype in one biological replicate. Image files were renamed with random codes by a person not involved in the scoring, so that genotype was concealed and image order was randomised. A single EM-experienced observer, blinded to genotype, then classified each capsid on the basis of tegument texture, thickness and density as showing either “normal tegument” (wild-type-like) or “condensed tegument” (hypo-P-like). Capsids that could not be assigned unambiguously, for example because of unfavourable orientation, oblique sectioning, mechanical damage or insufficient image quality, were recorded as “unclear”. Codes were broken only after all capsids had been scored, and no capsid was re-scored after unblinding.

### Software

Quantitative data were organized and preprocessed in Microsoft Excel. Curve fitting was performed in OriginPro (version 2025b, OriginLab). Statistical analyses and data visualization were performed in Python (version 3.12.3) with statsmodels (version 0.14), SciPy (v1.17.1), and matplotlib (version 3.10.8). Image acquisition, processing and fluorescence intensity measurements were performed in Fiji/ImageJ (v1.54, NIH)^44^, NIS-Elements AR (v5.42, Nikon), NIS-Elements AR Analysis (version 6.10.02, Nikon), and ZEN 3.0 (blue edition, Zeiss). Intrinsic disorder was predicted with IUPred3^29^. Proteomic data were processed with MSFragger (version 4.4.1) in FragPipe (version 24.0)^42^. Snapgene (v8.2.2) was used for primer design and sequence analysis. Final figure layouts were assembled in Affinity Designer (version 3.2.0, Canva).

### Statistics and reproducibility

No data were excluded from the analyses, except that, in the disorder enrichment analysis, proteins contributing fewer than two background crosslinks were excluded, because a per-protein background disorder distribution cannot be meaningfully defined from a single site. This criterion was pre-established and applied uniformly to all protein groups before comparison. Additionally, capsid images that could not be classified unambiguously in the blinded TEM assessment, as described above, were excluded from the statistical analysis. Investigators were blinded to genotype during the TEM capsid classification; all other analyses were performed unblinded. Throughout, biological replicates are independent infections, transfections, or nucleofections, and n is defined in each module together with the experimental unit. Significance is reported as ***P < 0.001, **P < 0.01, *P < 0.05, and ns, not significant; exact P values are given in the figure legends and in the Source Data file.

### Use of large language models

Large language models (ChatGPT, OpenAI; Claude, Anthropic) were used during preparation of this manuscript for two purposes: editing and restructuring of author-written text, and assistance with writing the analysis and plotting code used to generate figures from the underlying experimental data. All figures were generated from experimental data by code that was reviewed and validated by the authors. No figure, image, or micrograph was generated or modified by a generative artificial intelligence tool.

## Data availability

The mass spectrometry proteomics data have been deposited to the ProteomeXchange Consortium via the PRIDE partner repository under accession PXD081134 (reviewer token: i7QtoVmOJU3E)^45^. Source data underlying all graphs are provided as a Source Data file. The raw microscopy images generated during this study have been deposited to the BioImage Archive under the accession number S-BIAD3918. The dataset is currently under embargo and will be made publicly available upon publication.

## Code availability

The code used to analyze the local disorder at pp150 contact sites is available on GitHub at https://github.com/QuantitativeVirology/Contact-Disorder-Analysis. Taking the HCMV virion crosslinking dataset from Bogdanow et al. (2023) as input, it predicts local intrinsic disorder (IUPred3) at the pp150 contact sites of tegument, capsid, and host proteins and statistically compares them. Disorder prediction requires IUPred3, which must be obtained separately under its own license.

## Acknowledgements

This work was supported in part by a doctoral scholarship from the Studienstiftung des deutschen Volkes to YJ. Work in the Bosse lab was funded by the Deutsche Forschungsgemeinschaft (DFG, German Research Foundation) under Germany’s Excellence Strategy EXC 2155 project no. 390874280, the DFG-funded RTG 2771 Humans and Microbes, project number 453548970, the DFG-funded RTG 2887 VISION, project number 497350882, DFG-funded CRC 1648 Emerging Viruses project number SFB 1648/1 2024-512741711, by the Wellcome Trust through a Collaborative Award (209250/Z/17/Z), and the Leibniz ScienceCampus InterACt, funded by the BWFGB Hamburg and the Leibniz Association (W75/2022) InterACt and “Hamburg- X Infektionsforschung”. Moreover, the Bosse lab is funded through the DFG Research Unit FOR5200 DEEP-DV (443644894) project BO 4158/5-1 and BO 4158/5-2, DFG Research Unit FOR5898 AdBHealth (548065690) project BO 4158/9-1 and the German Center for Infection Research (DZIF) grants IICH TTU07.918, TTU 07.861 and 07.863. Additionally, this work was funded by DFG grant WI2043/4-1 to LW.

## Author contributions

Conceptualization: YJ, LW, JBB; Methodology: YJ, BB, JVE, LW; Investigation: YJ, LCR, BB, IG, BV, EC, LM, JVE; Formal Analysis: YJ, BB, JVE, LW, JBB; Resources: BB, JVE, LW, JBB; Visualization: YJ, LCR, JVE; Data Curation: YJ, BB; Funding Acquisition: BB, JVE, LW, JBB; Project Administration: LW, JBB; Supervision: BB, JVE, LW, JBB; Writing - original draft: YJ; Writing - review & editing: YJ, BB, JVE, LW, JBB.

## Supplementary Figures

**Supplementary Fig. 1:**
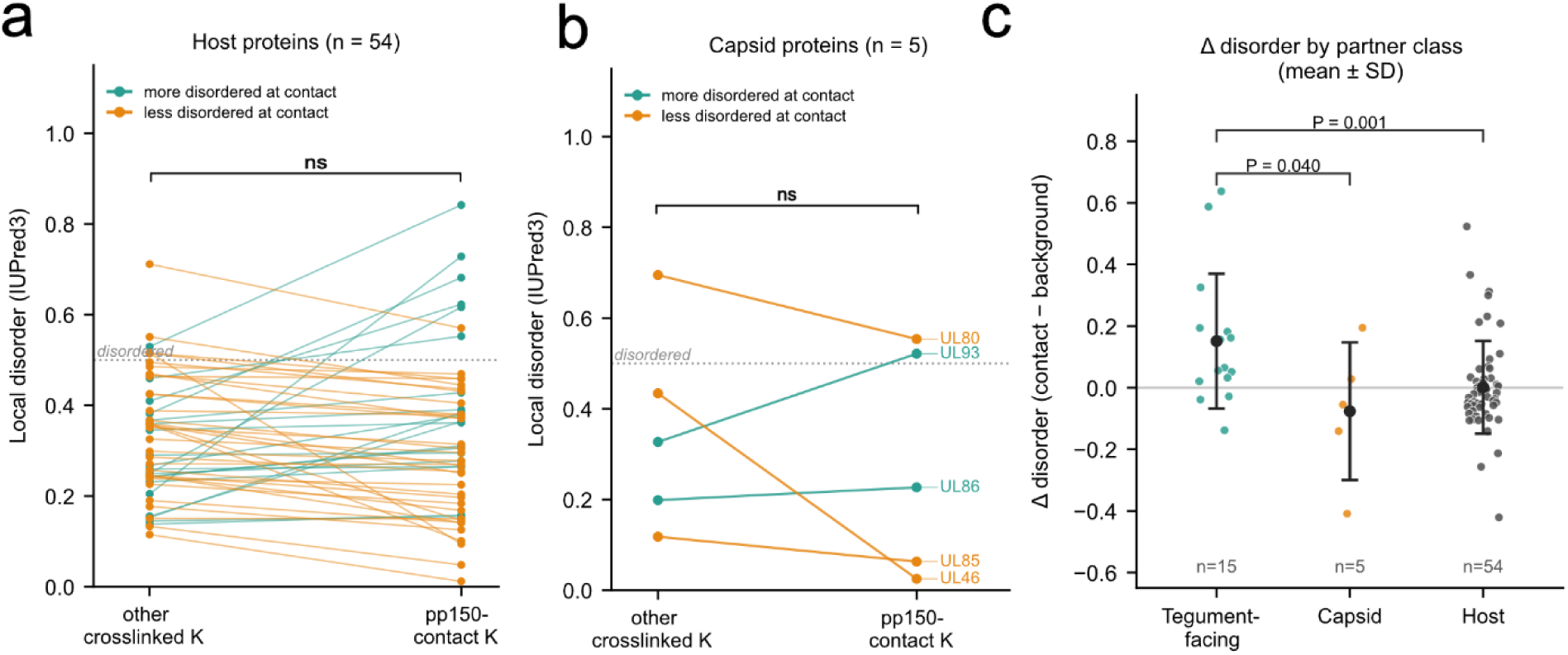
pp150 contacts tegument proteins through locally disordered regions, unlike its contacts with capsid or host proteins. Crosslinks involving pp150 were extracted from a published crosslinking mass spectrometry dataset of HCMV virions and mapped onto HCMV strain TB40-E sequences. An interface window was defined as ±7 residues centered on each lysine crosslinked to pp150, and its mean IUPred3 disorder was compared per protein with that of windows centered on the remaining crosslinked lysines of the same protein, which are equally solvent-exposed and reaction-competent and therefore provide an accessibility-matched background. **a**, Local disorder at pp150 contact sites in host proteins crosslinked to pp150, analysed exactly as in Fig. 2a. n = 54 host proteins (20 positive, 34 negative shifts). Mean Δdisorder = 0.0012 (standard deviation = 0.1505; 95% CI −0.0399 to 0.0423), median = - 0.0251; two-sided Wilcoxon signed-rank test, *W* = 597, *P* = 0.2103. **b**, Local disorder at pp150 contact sites in capsid and capsid-associated proteins, analysed exactly as in Fig. 2a. n = 5 proteins (2 positive, 3 negative shifts). Mean Δdisorder = −0.0764 (standard deviation = 0.2234; 95% CI −0.3538 to 0.2009), median = −0.0549; two-sided Wilcoxon signed-rank test, *W* = 5, *P* = 0.625. **c**, Dot plot comparing the enrichment of disorder at pp150 contact sites across the tegument, capsid and host groups. Black dots, mean; error bars, standard deviation. Groups were compared by a one-sided Mann–Whitney U test (tegument-facing versus capsid, *U* = 58, *P* = 0.0403; tegument-facing versus host, *U* = 611, *P* = 0.001).

**Supplementary Fig. 2:**
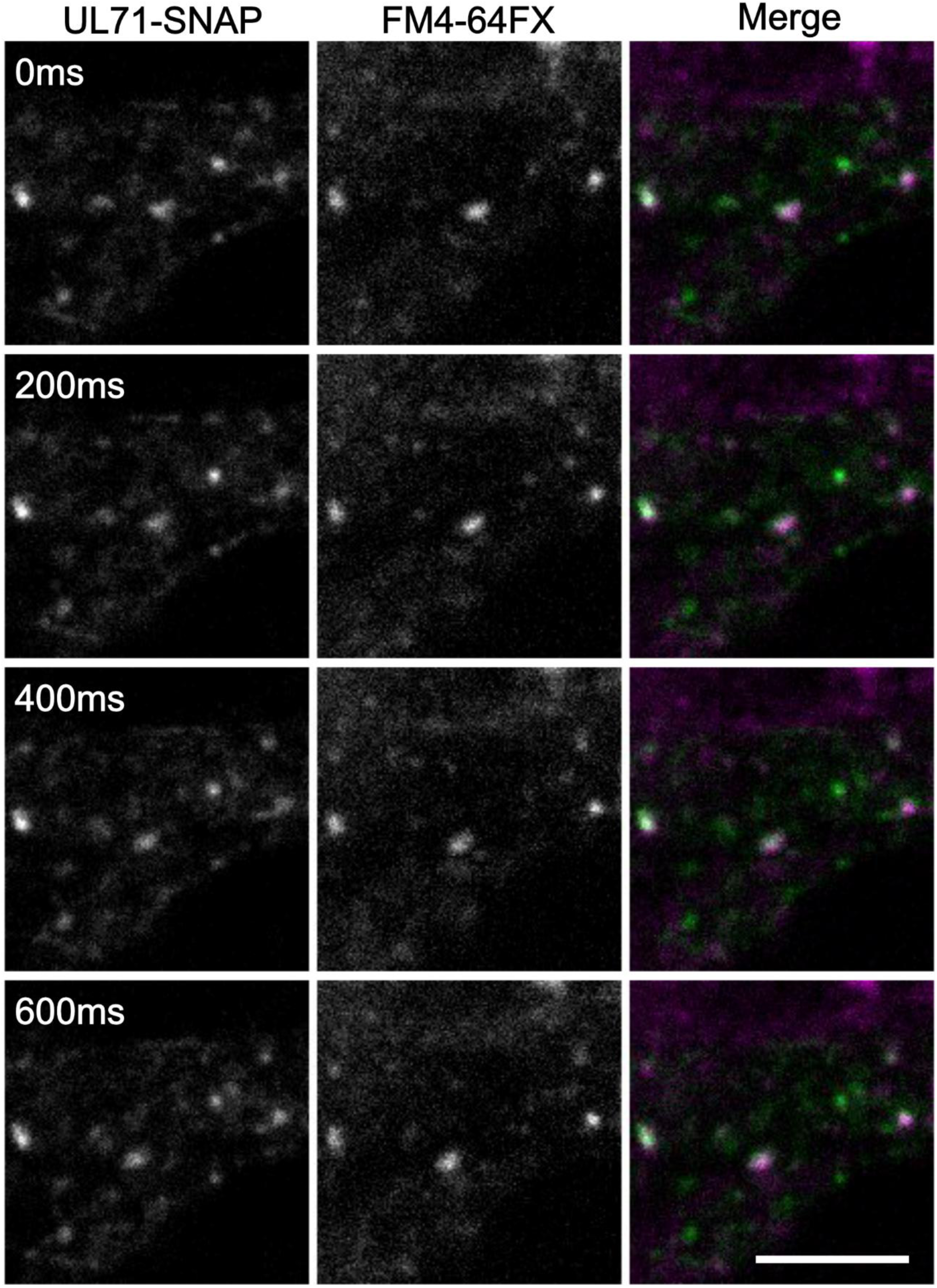
UL71-positive motile structures colocalize with the endosomal tracer FM4-64FX. Live-cell images of VeroB4 cells expressing UL71-SNAP, labelled with SNAP-Cell 505-STAR (green) and subsequently incubated with the membrane dye FM4-64FX (magenta), which initially labels the plasma membrane and is subsequently internalized by endocytosis, thereby marking endosomal compartments. Imaging 5 min after FM4-64FX addition revealed numerous small, UL71-positive puncta distributed throughout the cytoplasm, many of which exhibited rapid, directed movement consistent with vesicular transport. Motile UL71-positive puncta colocalized with FM4-64FX, consistent with an endosomal identity. Scale bar, 5 µm. Images are representative of 21 imaged cells from one transfection.

**Supplementary Fig. 3:**
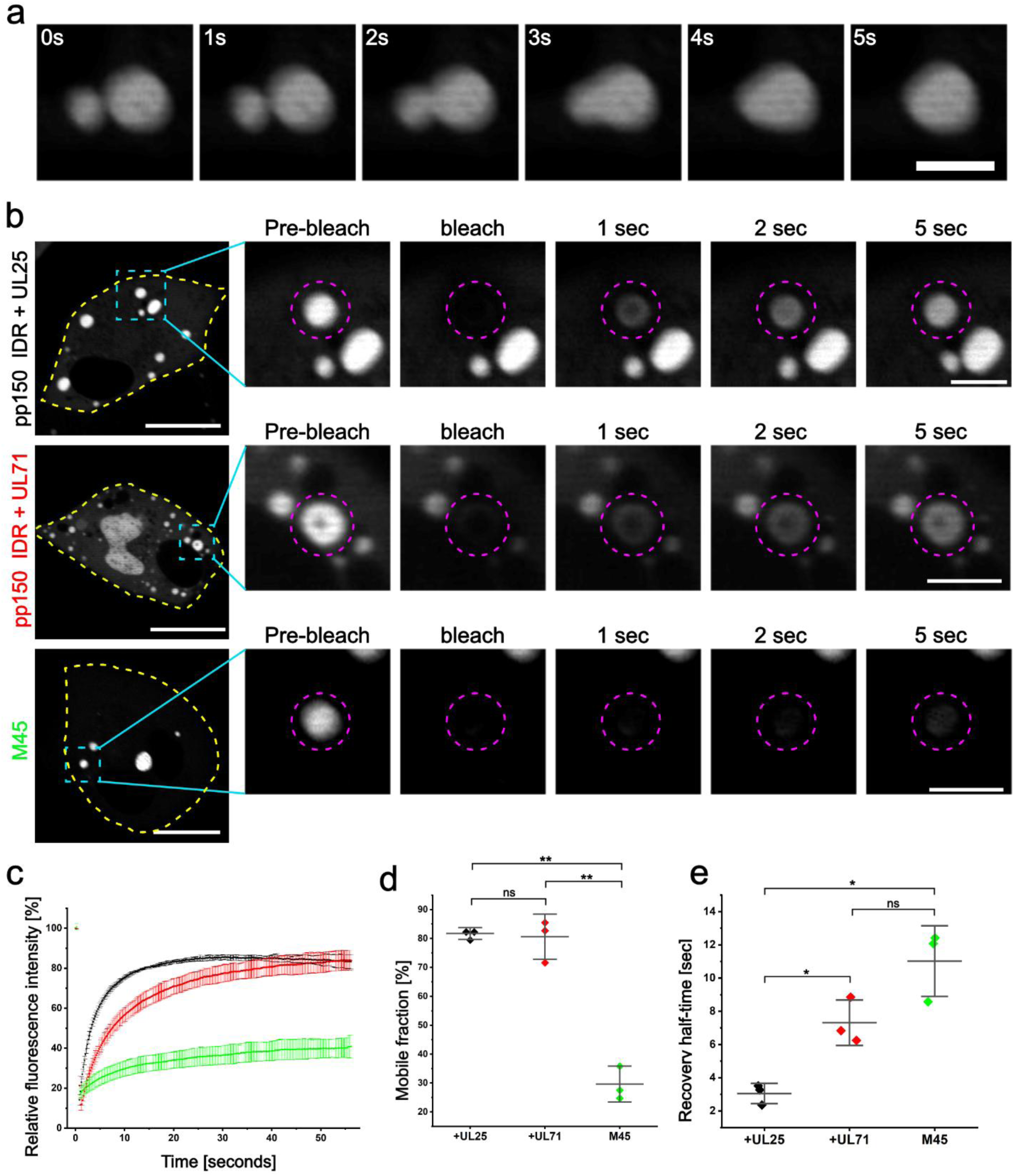
Co-condensates of the pp150 IDR with UL71 and UL25 exhibit liquid-like properties. VeroB4 cells co-expressing mScarlet3-pp150 IDR with either UL25-SNAP or UL71-SNAP were imaged live. mCherry-tagged M45 of murine cytomegalovirus, which forms stable, non-liquid aggregates, served as a negative control. **a**, Representative time series showing fusion of pp150-UL25 condensates. Scale bar, 5 µm. **b**, Representative fluorescence recovery after photobleaching (FRAP) experiment showing the whole cell (yellow outline), a magnified view of the condensate (cyan square) and the bleached region (magenta circle). Scale bars, 20 µm (overview) and 5 µm (inset). **c**, Quantification of fluorescence recovery over time for pp150 co-condensates with UL25-SNAP (black) or UL71-SNAP (red) and for mCherry-tagged M45 (green). Points, mean of three independent transfections (n = 3, 10–11 cells per experiment); error bars, standard deviation. **d**, Mobile fractions derived from fits of the recovery curves in c. Statistical comparisons with an unpaired two-sided Welch’s t-test (+UL25 versus M45, *P* = 2.32 × 10^-3^; +UL71 versus M45, *P* = 1.12 × 10^-3^; +UL25 versus +UL71, *P* = 8.28 × 10^-1^) **e**, Recovery half-times derived from the same fits. Statistical comparisons with an unpaired two-sided Welch’s t-test (+UL25 versus M45, *P* = 1.69 × 10^-2^; +UL71 versus M45, *P* = 7.45 × 10^-2^; +UL25 versus +UL71, *P* = 1.94 × 10^-2^) In d,e, dots represent individual experiments and bars show the mean ± standard deviation.

**Supplementary Fig. 4:**
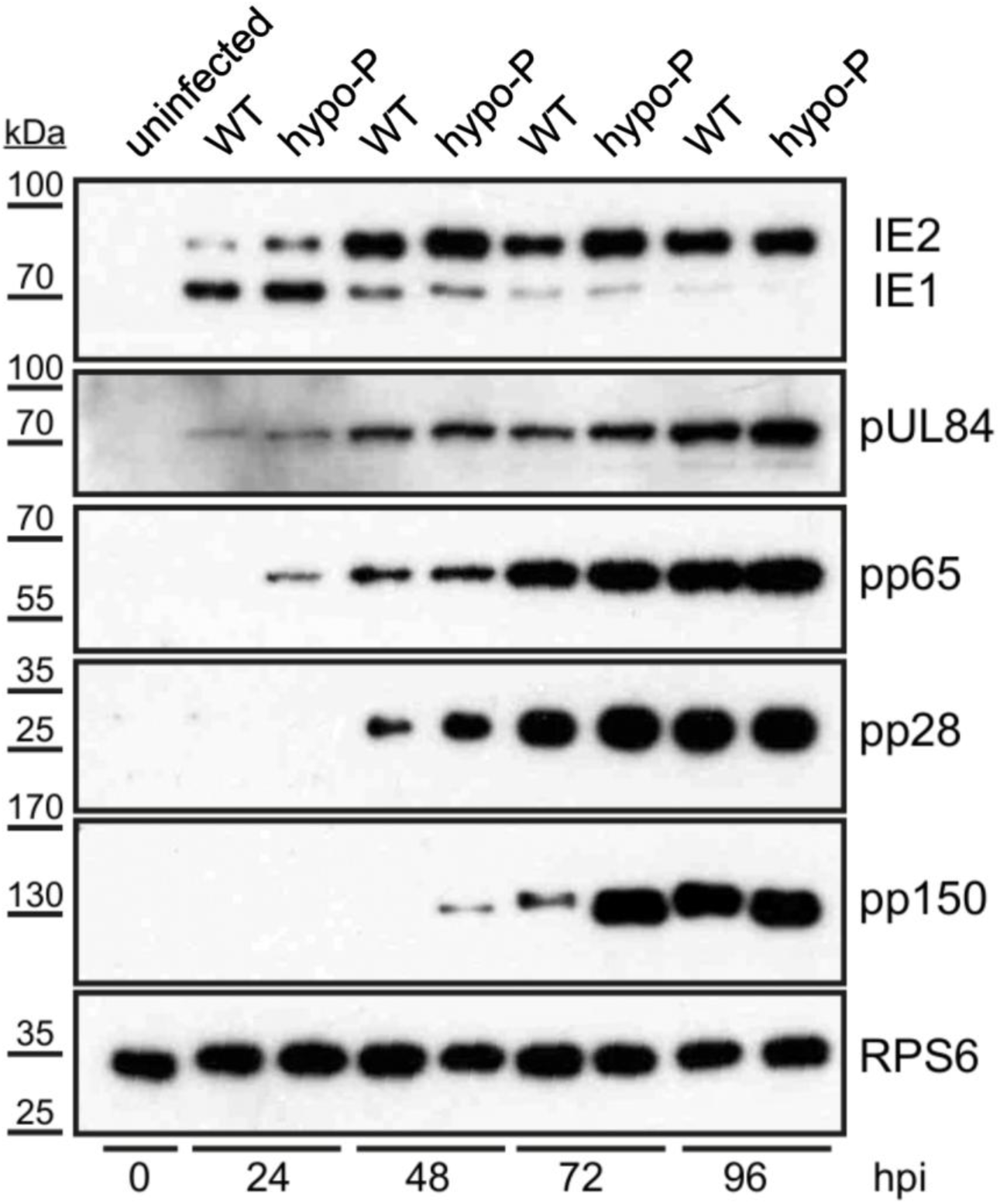
The hypo-P mutation leaves viral gene expression intact. Immunoblot analysis of HCMV protein accumulation in cells infected with wild-type virus or the hypo-P mutant over a 96 h time course (MOI = 5). Blots were probed for pp150 and for representative immediate-early, early and late viral proteins (IE1, IE2, UL84, pp65 and UL99). Comparable accumulation and temporal expression patterns in wild-type and hypo-P infection indicate that the growth defect of the hypo-P mutant is not caused by a broad impairment of viral gene expression or by loss of major tegument protein expression. RPS6 as loading control.

**Supplementary Fig. 5:**
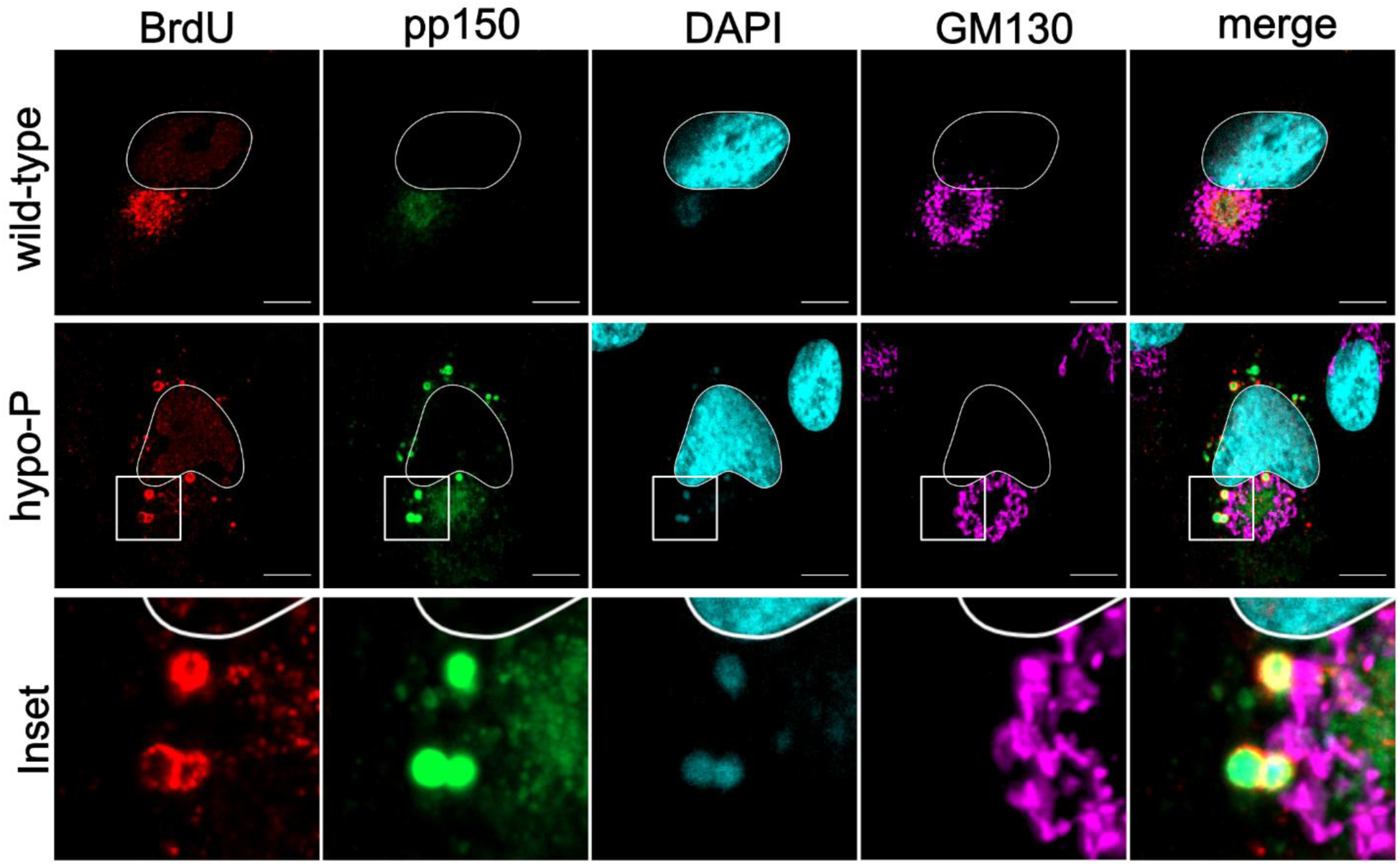
pp150-associated cytoplasmic aggregates contain viral DNA. Indirect immunofluorescence of HFFs infected with the indicated viruses at an MOI of 1 and fixed at 120 hpi. Nucleocapsids were detected by pulse labelling of newly synthesized viral genomes with 5-bromo-2′-deoxyuridine (BrdU), applied from 96 hpi for 20 h and followed by a 4 h chase (red). pp150, green; the cytoplasmic viral assembly compartment was stained with the cis-Golgi marker GM130, magenta; nuclei were stained with DAPI, blue, and outlined in white. Scale bars, 10 µm. Images are representative of 2 independent experiments.

**Supplementary Fig. 6:**
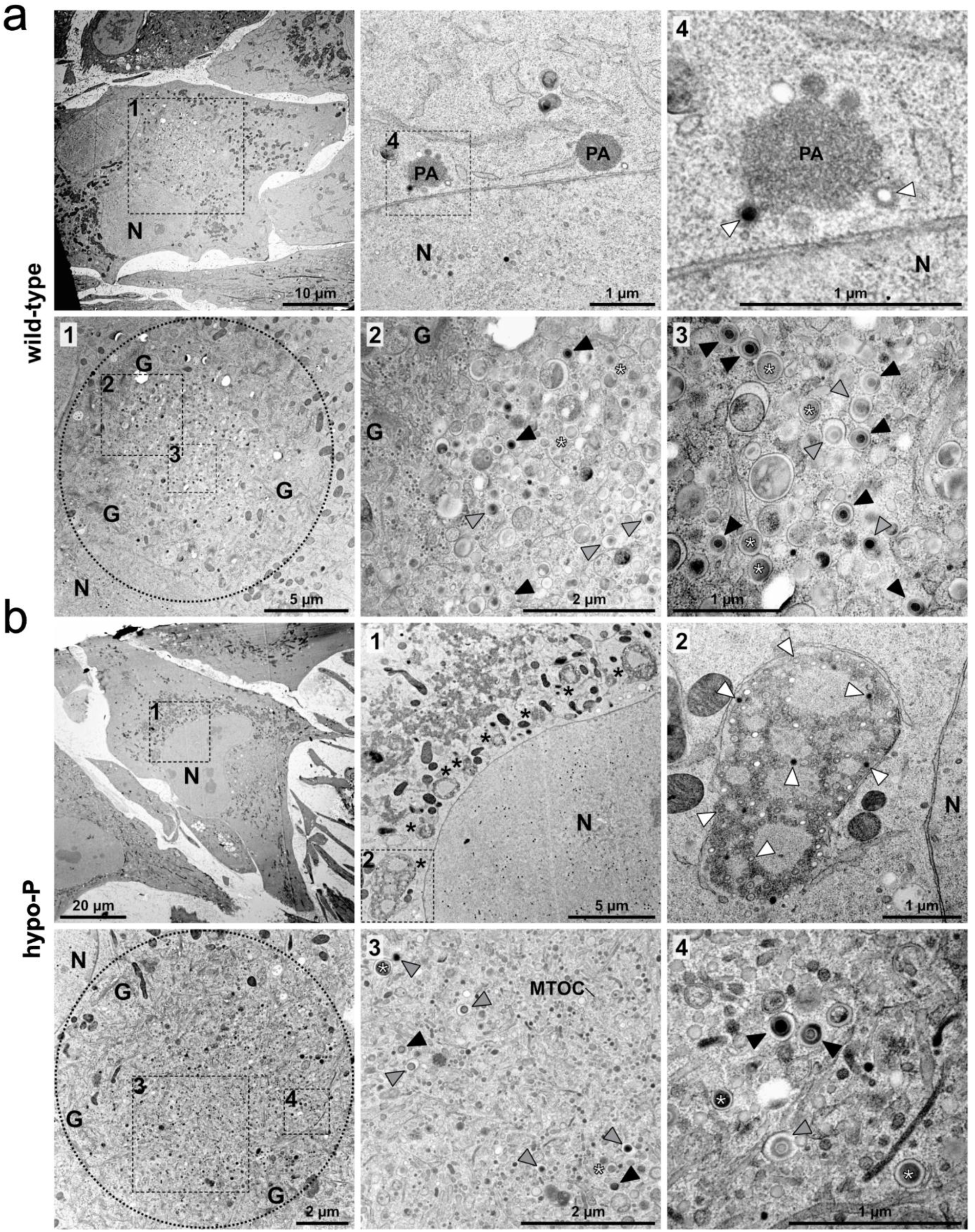
Ultrastructure of the cytoplasmic assembly compartment in wild-type and hypo-P infected cells. Electron micrographs of **a**, wild-type and **b**, hypo-P infected HFFs at 120 hpi. For each virus, an overview of the cytoplasmic viral assembly compartment (cVAC, dashed circle) and higher magnifications of the selected areas (dashed boxes, numbered) are shown. Capsids are labelled according to their stage of envelopment: free capsids, white arrowheads; budding capsids, grey arrowheads; enveloped capsids, black arrowheads. N, nucleus; G, Golgi; MTOC, microtubule-organizing center; PA, protein aggregates; black asterisks, capsid aggregates; white asterisks, dense bodies. Scale bars as indicated.

